# Loss of mitochondrial enzyme GPT2 leads to reprogramming of synaptic glutamate metabolism

**DOI:** 10.1101/2024.11.06.622291

**Authors:** Ozan Baytas, Shawn M. Davidson, Julie A. Kauer, Eric M. Morrow

## Abstract

Recessive loss-of-function mutations in the mitochondrial enzyme Glutamate Pyruvate Transaminase 2 (GPT2) cause intellectual disability in children. Given this cognitive disorder, and because glutamate metabolism is tightly regulated to sustain excitatory neurotransmission, here we investigate the role of GPT2 in synaptic function. GPT2 catalyzes a reversible reaction interconverting glutamate and pyruvate with alanine and alpha-ketoglutarate, a TCA cycle intermediate; thereby, GPT2 may play an important role in linking mitochondrial tricarboxylic acid (TCA) cycle with synaptic transmission. In mouse brain, we find that GPT2 is enriched in mitochondria of synaptosomes (isolated synaptic terminals). Loss of Gpt2 in mouse appears to lead to reprogramming of glutamate and glutamine metabolism, and to decreased glutamatergic synaptic transmission. Whole-cell patch-clamp recordings in pyramidal neurons of CA1 hippocampal slices from *Gpt2-*null mice reveal decreased excitatory post-synaptic currents (mEPSCs) without changes in mEPSC frequency, or importantly, changes in inhibitory post-synaptic currents (mIPSCs). Additional evidence of defective glutamate release included reduced levels of glutamate released from *Gpt2-*null synaptosomes measured biochemically. Glutamate release from synaptosomes was rescued to wild-type levels by alpha-ketoglutarate supplementation. Additionally, we observed evidence of altered metabolism in isolated *Gpt2-*null synaptosomes: decreased TCA cycle intermediates, and increased glutamate dehydrogenase activity. Notably, alterations in the TCA cycle and the glutamine pool were alleviated by alpha-ketoglutarate supplementation. In conclusion, our data support a model whereby GPT2 mitochondrial activity may contribute to glutamate availability in pre-synaptic terminals, thereby highlighting potential interactions between pre-synaptic mitochondrial metabolism and synaptic transmission.

## Introduction

Glutamate is the main excitatory neurotransmitter in the central nervous system (CNS) [1, 2]. In addition, glutamate serves as a metabolite involved in a number of crucial cellular functions, including the tricarboxylic acid (TCA) cycle [3, 4], nitrogen balance [5, 6], protein synthesis [7], glutathione synthesis [8], and gamma-aminobutyric acid (GABA) synthesis, the major inhibitory neurotransmitter in the brain [9]. Glutamate metabolism is interwoven with synaptic transmission and various pathways maintain metabolic supply at the synapse. Notably, the glutamate-glutamine cycle has been proposed to ensure continuous glutamate availability for excitatory neurotransmission; however accumulating evidence suggests that glutamate metabolism may interact with the tricarboxylic acid (TCA) cycle to support energetics in both astrocytes and neurons via transamination or deamination [10].

Human genetic mutations offer a powerful opportunity to identify metabolic pathways required for cognition. Previously, we reported autosomal recessive, loss-of-function mutations in the mitochondrial enzyme Glutamate Pyruvate Transaminase 2 (*GPT2*) that lead to a neurodevelopmental disorder involving cognitive disability [11]. GPT2 catalyzes a reversible transamination, transferring an amino group from glutamate to pyruvate synthesizing alanine and alpha-ketoglutarate, a TCA cycle intermediate. GPT2 is localized to the mitochondria while its paralog, GPT1 catalyzes the same reaction in the cytoplasm [11, 12]. GPT2 has been studied in the context of cancer wherein it has been reported to play an essential role in cell growth through replenishment of TCA cycle intermediates, a process known as anaplerosis [13–16]. The function of GPT2 in the nervous system is not well understood. GPT2 is enriched in the brain and upregulated during postnatal brain development, a time of prominent synaptogenesis [11, 17]. The contributions of GPT2 to synaptic transmission are unknown.

In this study, we report that GPT2 is enriched in the mitochondria of synaptosomes (isolated synaptic terminals). We also provide evidence to indicate that loss of GPT2 may lead to decreased glutamate availability for synaptic transmission, as well as altered glutamate and glutamine metabolism in synaptosomes. GPT2 produces glutamate with alanine and alpha-ketoglutarate as precursors, and through electrophysiology, we find that glutamatergic transmission in mouse brain slices is impaired. Also, using biochemical techniques, we observed reduced glutamate levels released from *Gpt2-*null synaptosomes. In addition, we find evidence for altered glutamate and glutamine metabolism in isolated synaptic terminals, including increases in glutamine levels, and increased activity of glutamate dehydrogenase. Importantly, we demonstrate that alpha-ketoglutarate was able to raise glutamate levels and correct measures of altered glutamine metabolism in *Gpt2-*null synaptosomes. Finally, we find deficits in the TCA cycle, such as reduced TCA cycle intermediates, in *Gpt2-*null synaptosomes, although glutamine entry into the TCA cycle remains intact and indicators of energetics appear largely unchanged. Reduced levels of TCA cycle intermediates is again rescued by alpha-ketoglutarate supplementation in *Gpt2-*null synaptosomes. Overall, our results suggest that GPT2 plays a role in sustaining neurotransmitter availability, and in the absence of GPT2, glutamatergic synaptic transmission and energetics appear to be partly compensated by reprogrammed glutamate metabolism which may be ameliorated by alpha-ketoglutarate supplementation.

## Materials and Methods

### Ethics Statement

All experiments that involved mice were done in accordance with the National Institutes of Health *Guide for the Care and Use of Laboratory Animals* [18] and approved by the Brown University Institutional Animal Care and Use Committee.

### Animals

*Gpt2-*null animals were obtained from Knockout Mouse Project at University of California, Davis (RRID: MMRRC_047980-UCD), genotyped and maintained as described previously [11]. The background of all mice was C57BL6/J (Jackson Laboratory Strain No. 664, RRID: IMSR_JAX:000664). The mice were maintained under a 12-hour light/dark cycle (lights on at 7 am and off at 7 pm). The mice had ad libitum access to feed and water. All experiments were conducted on mice at postnatal day 18 (P18).

### Synaptosome Preparation

The protocol was adapted from [19] and [20]. Immediately after cervical dislocation, the brain was removed at an angle with double-ended surgical spatula so that the midbrain, pons, medulla, and cerebellum were excluded (Fig. 1A). Olfactory bulb was also excluded. The tissue was rinsed in ice-cold 1X brain homogenization buffer (0.32 M sucrose, 1 mM EDTA, 5 mM Tris, pH 7.4). Fresh 1X brain homogenization buffer (10 mL/g) was added, and the brain was cut into small pieces with surgical scissors. A Kimble pestle was used to homogenize the tissue with 12 strokes (each 4 sec). The tube was spun at 1300 x *g* for 3 min at 4°C. The first supernatant was stored on ice and the pellet was re-suspended in half starting volume with 1X brain homogenization buffer. The pellet was homogenized with a Kimble pestle with 8 strokes (each 4 sec). The tubes were spun at 1300 x *g* for 3 min at 4°C. The first and second supernatants were combined, and the sample was spun at 10000 x *g* for 10 min at 4°C with a Sorvall RC6Plus centrifuge using the SS34 rotor. The pellet was re-suspended in 10 mL per gram of original tissue with 15% Percoll (Percoll, Cytiva 17089102, prepared in 1X brain homogenization buffer). The suspension was carefully layered onto a Percoll gradient (23%-40%) and the tube was spun at 22000 x *g* for 10 min at 4°C. The top layer contains myelin and associated membranes, the middle layer contains synaptosomes and associated membranes and the bottom layer contains the mitochondrial fraction. After carefully removing the upper layer, the synaptosomes were aspirated and re-spun with 4 volumes of 1X brain homogenization buffer, at 16000 x *g* for 10 min at 4°C. The supernatant was carefully removed, and 4 volumes of Krebs-like buffer (118 mM NaCl, 5 mM KCl, 1 mM MgCl_2_, 1.2 mM CaCl_2_, 0.1 mM Na_2_HPO_4_.H_2_O, 20 mM HEPES, 10 mM glucose, pH 7.4) was added. The tube was spun at 18000 x *g* for 10 min at 4°C. The final pellet was flash frozen in liquid nitrogen for experiments in which baseline metabolite levels were determined. For all other experiments, the final pellet was re-suspended in the Krebs-like buffer with 0.5 mM glutamine (Invitrogen, 25030081).

**Figure 1.**
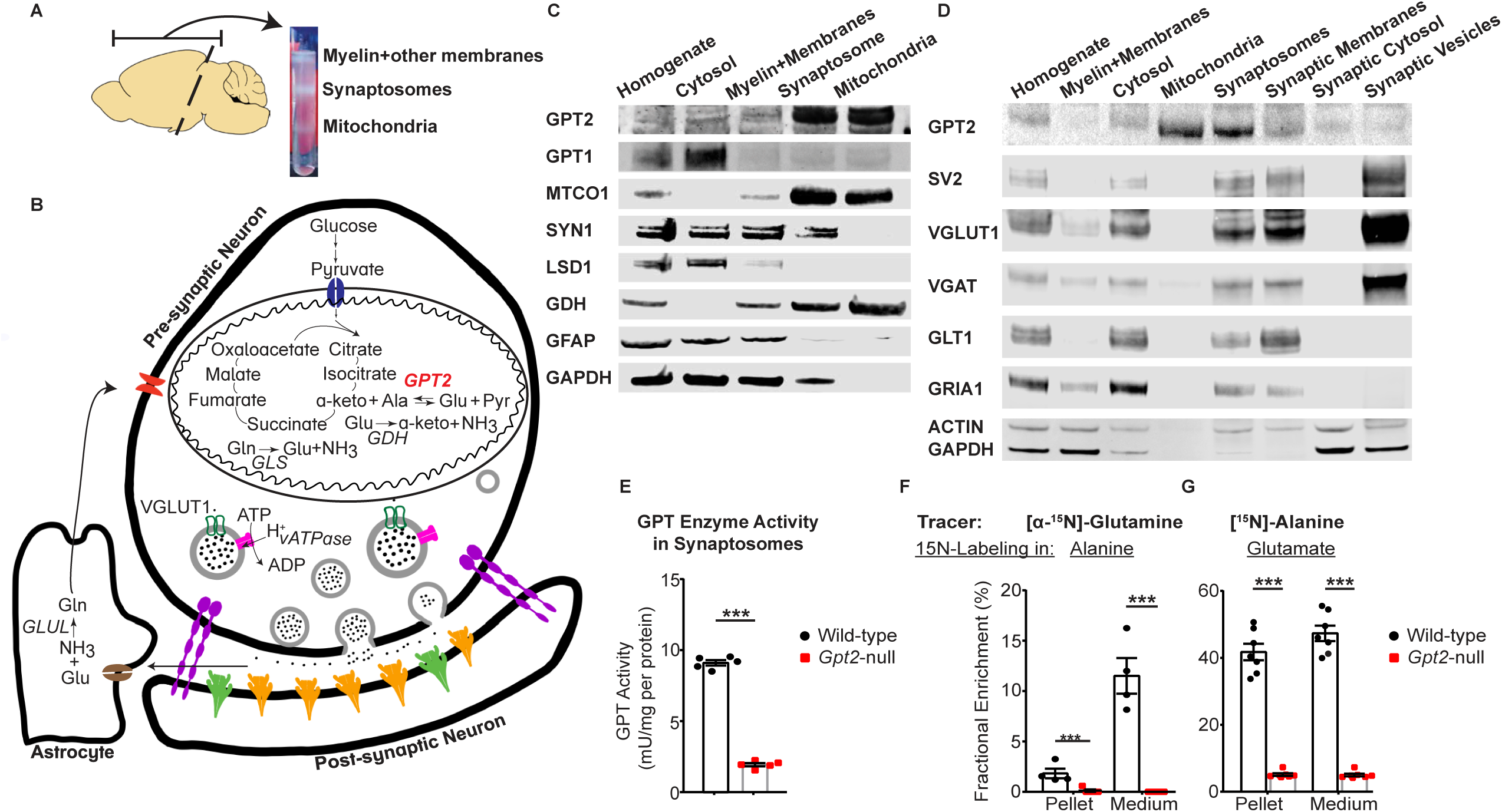
GPT2 is the primary glutamate pyruvate transaminase in isolated synaptic terminals. **A.** Fractionation of the mouse brain into fractions containing myelin and other membranes, synaptosomes (isolated synaptic terminals) and mitochondria. Note that the synaptosomes are extracted from brain tissue containing only the forebrain, excluding olfactory bulb, midbrain and the brain stem. **B.** Metabolic pathways relevant to GPT2 and glutamate metabolism in synaptosomes including pre-synaptic neuron, post-synaptic neuron and astrocytic components. Ala: alanine, a-keto: alpha-ketoglutarate, Glu: glutamate, Gln: glutamine, Pyr: pyruvate, *GDH*: glutamate dehydrogenase, *GLS*: glutaminase, *GLUL*: glutamine synthetase. **C.** GPT2 is enriched in synaptosomes and mitochondria. Western blotting of the homogenate, cytosolic, myelin and membranes, synaptosomes and mitochondria fractions obtained from a wild-type mouse at postnatal day 18 (P18) for GPT2, GPT1, MTCO1 (mitochondrial marker, subunit 1 of complex IV cytochrome c oxidase), SYN1 (synapsin I), LSD1 (nuclear marker, lysine-specific demethylase 1), GDH (glutamate dehydrogenase), GFAP (glial fibrillary acidic protein) and GAPDH (cytosolic marker, glyceraldehyde 3-phosphate dehydrogenase). **D.** GPT2 is enriched in mitochondria of synaptosomes. Western blotting of synaptosomes that were fractionated further into synaptic membrane, synaptic cytosol and synaptic vesicles obtained from a wild-type mouse at P18 for GPT2, SV2 (synaptic vesicle protein 2), VGLUT1 (vesicular glutamate transporter 1), VGAT (vesicular GABA transporter), GLT1 (glutamate transporter 1, astrocytic), GRIA1 (AMPA receptor subunit 1), ACTIN, GAPDH. **E.** Overall GPT enzyme activity in *Gpt2-*null is greatly reduced in *Gpt2-*null synaptosomes at P18. Each dot represents the total enzymatic activities of GPT2 and GPT1 in synaptosome samples obtained from different wild-type (black) or *Gpt2-*null (red) mice. The enzymatic activities are expressed as mU/mg per protein. *P* <0.0001. **F.** Glutamine cannot be used as a precursor for alanine in *Gpt2-*null synaptosomes at P18. Isotope tracing in wild-type (black) and *Gpt2-*null (red) synaptosomes at P18 using glutamine labeled with heavy nitrogen at the amine position. Each dot represents fractional enrichment of labeled pool of alanine in synaptosome from a different animal, expressed as percentage of total alanine pool. For pellet, \*\**P* = 0.006 and for medium, \*\*\**P* = 0.00016. **G.** Alanine and alpha-ketoglutarate can be used as precursors for glutamate in wild-type but not in *Gpt2-*null synaptosomes at P18. Isotope tracing in wild-type (black) and *Gpt2-* null (red) synaptosomes at P18 using heavy nitrogen labeled alanine. Each dot represents fractional enrichment of labeled pool of glutamate in synaptosome from a different animal, expressed as percentage of total glutamate pool. For pellet and medium, \*\*\**P* <0.0001.

For isolation of synaptic vesicles, after the synaptosomes were washed with 4 volumes of Krebs-like buffer, the pellet was re-suspended in 450 µl ice-cold ddH_2_O. The pellet was homogenized by pipetting. 4.95 µl 1M HEPES (pH7.4, with NaOH) was immediately added and the mixture was incubated on ice for 30 min. The mixture was spun at 43,100 x *g* with SS-34 rotor at 4°C for 15 min. The supernatant was distributed equally to TLA 120.1 (Beckman Coulter) rotor thick-walled polycarbonate tubes (Beckman Coulter, 343776). The tubes were spun at 165,000 x *g* at 4°C for 1 hour. The pellet was re-suspended in 30 µl RIPA buffer (50 mM Tris, 0.15 M NaCl, 1 mM EDTA, 1% TritonX-100, 0.5% (w/v) sodium deoxycholate, 0.1% SDS) with 1X PhosStop and 1X PIC for Western blotting.

### Western Blotting

Tissues were homogenized in RIPA buffer (50 mM Tris, 0.15 M NaCl, 1 mM EDTA, 1% TritonX-100, 0.5% (w/v) sodium deoxycholate, 0.1% SDS) with 1X PhosStop and 1X PIC. Protein samples (20 µg) were incubated in 1X NuPage Sample buffer (Invitrogen, NP0007) and 1X NuPage reducing agent (Invitrogen, NP0004) at 70°C for 10 min and immediately placed on ice. The samples were run in NuPage 4-12% Bis-Tris gel (Invitrogen, NP0321) at 170 V until the dye front was at the end of the gel. The gel was transferred to a nitrocellulose membrane (Invitrogen, LC2000) in 1X FisherScientific Pierce Western Blot Transfer Buffer (35040) with 20% methanol at 30 V for 1 hour. The blot was blocked with Li-Cor Blocking Buffer (Li-Cor Biosciences, 927-50000) for 30 min and permeabilized for 5 min in TBST (FisherScientific, BP2411) with 0.05% Tween20 (Sigma-Aldrich, P7949). The blot was incubated with primary antibodies in Li-Cor Intercept Antibody Diluent (Li-cor Biosciences, 927-65001) at 4°C overnight. The blot was washed 3 x 5 min with TBST, incubated with secondary antibodies diluted in TBST for 1 hour, washed with TBST and finally placed in TBS. The blot was imaged using the Li-Cor Odyssey CLx Imaging System (resolution: 84 µm, background subtraction: median) and analyzed using Image Studio Lite software (RRID:SCR_013715). Protein amount in each sample was determined using the bicinchoninic acid (BCA) assay (ThermoScientific Pierce, PI23227). Primary antibodies used in the study: mouse anti-ACTIN (Sigma, A3853, RRID:AB_262137, 42 kDa, 1:2000), mouse anti-GAPDH (Sigma, G8795, RRID:AB_1078991, 37 kDa, 1:40,000), rabbit anti-GDH (Invitrogen, PA5-29492, RRID:AB_2546968, 55 kDa, 1:2000), rabbit anti-GLS1 (Proteintech, 20170-1-AP, RRID:AB_10665373, 66 kDa, 1:500), mouse anti-GLT1 (Millipore, MAB2262, RRID:AB_10615610, 62 kDa, 1:1000), mouse anti-GLUL (BD Biosciences, 610517, RRID:AB_397879, 42 kDa, 1:2000), mouse anti-GOT1 (Invitrogen, MA5-36093, RRID:AB_2890416, 45 kDa, 1:500), rabbit anti-GOT2 (Sigma, HPA018139, RRID:AB_1849903, 45 kDa, 1:1000), mouse anti-GPT1 (Sigma, SAB1412234, RRID:AB_2885183, 55 kDa, 1:1000), rabbit anti-GPT2 (Proteintech, 16757-1-AP, RRID:AB_2112098, 58 kDa, 1:600), rabbit anti-GRIA1 (abcam, ab31232, RRID:AB_2113447, 100 kDa, 1:1000), rabbit anti-LSD1 (CellSignaling, 2184, RRID:AB_2070132, 110 kDa, 1:1000), mouse anti-MTCO1 (abcam, ab14705, RRID:AB_2084810, 40 kDa, 1:2000), mouse anti-SV2 (DSHB, SV2, RRID:AB_2315387, 90 kDa, 1:1000), rabbit anti-SYNAPSINI (Millipore, AB1543P, RRID:AB_90757, 77&80 kDa, 1:1000), guinea pig anti-VGAT (SynapticSystems, 131004, RRID:AB_887873, 50 kDa, 1:1000), guinea pig anti-VGLUT1 (SynapticSystems, 135304, RRID:AB_887878, 50 kDa, 1:1000). Secondary antibodies used in the study: goat anti-Rabbit IRDye 680RD (Li-Cor Biosciences, 926-68071, RRID:AB_10956166, 1:20,000), goat anti-Rabbit IRDye 800CW (Li-Cor Biosciences, 925-32211, RRID:AB_2651127, 1:20,000), goat anti-Mouse IRDye 680RD (Li-Cor Biosciences, 925-68070, RRID:AB_2651128, 1:20,000), goat anti-Mouse IRDye 800CW (Li-Cor Biosciences, 926-32210, RRID:AB_621842, 1:20,000), donkey anti-Guinea pig IRDye 680RD (Li-Cor Biosciences, 925-68077, RRID:AB_2814914, 1:20,000), donkey anti-Guinea pig IRDye 800CW (Li-Cor Biosciences, 925-32411, RRID:AB_2814905, 1:20,000).

### GPT Enzyme Activity Assay

GPT activity assay was performed using the Alanine Aminotransferase (ALT) Assay Kit (Sigma-Aldrich, MAK052) according to the manufacturer’s manual. Synaptosomes were resuspended in 1X ALT Assay buffer. 5 µg protein sample was used. Protein amount in each sample was determined using the BCA assay (ThermoScientific Pierce, PI23227). BioTek Cytation5 plate reader was used to detect fluorescence; Gen5 software was used to analyze the data (RRID:SCR_017317).

### Metabolite Detection By Liquid Chromatography - Mass Spectrometry (LC-MS)

For all experiments in which metabolites were detected in synaptosomes without any additional supplements, synaptosomes were collected as above and the final pellet was flash frozen in liquid nitrogen. The pellet was homogenized in 400 µL of 80% (vol/vol) methanol cooled to −80°C with Kimble pestle and motor, then vortexed for 1 min, incubated at −80°C for 4 hours. The samples were spun at 14000 x *g* for 10 min at 4°C and the pellet was resuspended in 400 µL of 80% methanol while the supernatant was stored at −80°C. The suspension was incubated at −80°C for 30 min and then spun at 14000 x *g* for 10 min at 4°C. The supernatants were combined and re-spun at 14000 x *g* for 10 min at 4°C. The supernatants were concentrated using a Savant SpeedVac vacuum concentrator (Thermo Scientific) at ambient temperature in 1.5-mL Eppendorf tubes. Liquid chromatography with tandem mass spectrometry was done at Beth Israel Deaconess Medical Center Mass Spectrometry Facility. The samples were re-suspended in liquid chromatography (LC) / mass spectrometry (MS)-grade water and run-in tandem LC-MS/MS. Samples were injected into hydrophilic interaction liquid chromatography (HILIC) at high pH using HPLC coupled to a 5500 QTRAP mass spectrometer (AB/SCIEX). Selected Reaction Monitoring (SRM) mode for 300 transitions with positive/negative polarity switching fragmented precursor ions and selected for product ions. Peak areas for each detected metabolite were integrated using MultiQuant software (AB/SCIEX). All data were normalized by median normalization of at least 100 metabolite peaks. Unpaired Student’s t-test assuming equal variance was used to generate a p-value.

For all experiments that involve additional supplement with 1.6 mM alanine (Sigma, A7627) and 1.6 mM alpha-ketoglutarate (Sigma, 75890), synaptosomes in all conditions were incubated in Krebs-like buffer with 0.5 mM glutamine for 15 min at 37°C. For evoked glutamate release, a final concentration of 50 mM KCl was added, and the suspension was incubated for 1 min at 37°C. The pellet was spun immediately after at 16000 x *g* at 4°C for 1 min. LC−MS analysis for soluble metabolites was achieved on the Q Exactive PLUS hybrid quadrupole-orbitrap mass spectrometer (Thermo Scientific) coupled with hydrophilic interaction chromatography (HILIC). To perform the LC separation, an XBridge BEH Amide column (150 mm × 2.1 mm, 2.5 μm particle size, Waters, Milford, MA) was used with a gradient of solvent A (95%:5% H_2_O: acetonitrile with 20 mM ammonium acetate, 20 mM ammonium hydroxide, pH 9.4), and solvent B (100% acetonitrile). The gradient was 0 min, 85% B; 2 min, 85% B; 3 min, 80% B; 5 min, 80% B; 6 min, 75% B; 7 min, 75% B; 8 min, 70% B; 9 min, 70% B; 10 min, 50% B; 12 min, 50% B; 13 min, 25% B; 16 min, 25% B; 18 min, 0% B; 23 min, 0% B; 24 min, 85% B; 30 min, 85% B. The flow rate was 150 μL/min; the injection volume was 10 μL; the column temperature was 25°C. MS full scans were in negative ion mode with a resolution of 140,000 at m/z 200 and scan range of 75-1000 m/z. The automatic gain control (AGC) target was 1 × 10^6^.

#### Isotope Labeling in Synaptosomes

Synaptosome metabolite pool was labeled with 0.5 mM [α-^15^N]-glutamine (Sigma-Aldrich, 486809), 0.5 mM [U-^13^C]-glutamine (Sigma-Aldrich, 605166) or 1.6 mM [α-^15^N]-alanine (Cambridge Isotope Laboratories, NLM-454) for 15 min at 37°C. All conditions had labeled or unlabeled 0.5 mM glutamine in Krebs-like buffer.

### Brain Slice Preparation and Electrophysiological Recordings

Coronal slices (250 µm) were prepared from anesthetized mice at postnatal day 18, as described previously [21]. Briefly, the brain was immediately extracted and placed in a vibratome (Leica, VT1000). Slices were cut in ice-cold oxygenated artificial cerebrospinal fluid (ACSF, in mM): 119 NaCl, 2.5 KCl, 2.5 CaCl_2_.2H_2_O, 1.0 NaH_2_PO_4_.H_2_O, 1.3 MgSO_4_.7H_2_O, 26.0 NaHCO_3_, 11 glucose. Slices were recovered in the same oxygenated ACSF solution at room temperature for 1 hour and then transferred to a recording chamber where they were continuously submerged in ACSF at 28°C with a flow rate of 1-2 ml/min.

Whole cell recordings of CA1 pyramidal neurons were performed using a MultiClamp 700B Amplifier (Molecular Devices). Signals were low-pass filtered at 3 kHz and digitized at 10 kHz via a Digidata 1550 digitizer (Molecular Devices). Patch electrodes were fabricated from borosilicate glass capillaries (Sutter Instruments) using a P-97 micropipette puller (Sutter Instruments). Patch pipettes had a resistance of 3-8 MOhm. For determining intrinsic cell properties, the patch pipette internal solution consisted of (in mM): 125.0 KCl, 2.8 NaCl, 10.0 HEPES, 2.0 MgCl_2_, 2.37 ATP-Mg, 0.32 GTP-Na, 0.6 EGTA, (pH 7.23-7.28, 270-278 mOsm). Capacitance and membrane resistance were calculated from voltage response during injection of −100 pA.

For measuring miniature synaptic currents, the patch pipette internal solution consisted of (in mM): 125.0 CsCl, 2.8 NaCl, 10.0 HEPES, 2.0 MgCl_2_, 2.37 ATP-Mg, 0.32 GTP-Na, 0.6 EGTA, (pH 7.23-7.28, 270-278 mOsm). At least 80 events were detected in ASCF with 1 µM TTX (Tocris, 1078) and 33 µM Bicuculline (Tocris, 0130) for miniature excitatory synaptic currents or with 1 µM TTX and 10 µM DNQX (Sigma, D0540) for miniature inhibitory synaptic currents, while the cell was voltage-clamped at - 80mV. The series resistance was monitored throughout the recording and the cell was excluded from analysis if it changed by 20%. Miniature events with multiple consecutive peaks or “shoulders” were counted for frequency but excluded from amplitude calculations. For miniature excitatory currents only: the traces were low-pass (Gaussian) filtered with a −3dB cutoff frequency of 1000 Hz for better visualization of miniature events with smaller amplitudes. The miniature events were detected using a template compiled from each recording individually in Clampfit software (SCR_011323).

For paired pulse ratio analysis, the patch pipette internal solution consisted of (in mM): 125.0 KCl, 2.8 NaCl, 10.0 HEPES, 2.0 MgCl_2_, 2.37 ATP-Mg, 0.32 GTP-Na, 0.6 EGTA, 5 mM QX-314 (Tocris, 2313) (pH 7.23-7.28, 270-278 mOsm). The external solution consisted of ASCF with 50 µM d-APV (Tocris, 0106) and 100 µM picrotoxin (Sigma, P1675). CA3 was surgically removed from the slice to minimize epileptic activity. To stimulate Schaffer collaterals, bipolar stainless steel microelectrodes with 2-5 MOhm impedance (FHC, UESMEGSEKNNM) were gently placed approximately 200 µm away from the recorded pyramidal neuron. Paired pulses, 50 msec apart, were used to achieve a stable current response of approximately 150 - 300 pA. Paired pulse ratio was calculated by dividing the peak amplitude of the second response to that of the first. All electrophysiological recordings were analyzed using Clampfit software.

### Glutamate Measurement By Enzymatic Method

Free glutamate levels in the cytosolic fraction, that is the supernatant obtained along with the pellet that was subsequently re-suspended in Percoll (see *Synaptosome Preparation*), were determined by enzymatic method using glutamate dehydrogenase. Protein concentration was quantified by the bicinchoninic acid (BCA) assay (ThermoScientific Pierce, PI23227). 100 µg of each sample was added to their respective wells in a black clear-bottom 96-well plate (Greiner Bio-One, 655090) and the final volume was completed to 200 µl with Krebs-like buffer. NADP^+^ (Sigma, N5755) at a final concentration of 1 mM was added to each well. After 6 minutes of equilibration, five units of glutamate dehydrogenase (Sigma, G2626) was added to each well. The excitation and emission filters were 340 and 460 nm, respectively. All readings were taken at 37°C. Glutamate standard (Sigma, G1251) was used for calibration. To validate the viability of synaptosomes, 50 mM KCl was added to synaptosomes immediately after the fluorescence reached a plateau following glutamate dehydrogenase addition.

### Electron Microscopy

The animals were perfused with 1X PBS and then 2.5% glutaraldehyde, 2% paraformaldehyde, 2 mM calcium chloride in 0.15 M sodium cacodylate buffer. The 100 µm thick tissue sections were obtained using a vibratome and fixed in glutaraldehyde/paraformaldehyde/calcium chloride/sodium cacodylate buffer for 3 hours at 4°C. The sections were washed in cold 0.15 M sodium cacodylate with 2 mM calcium chloride for 3 minutes, 5 times and then in filtered ddH_2_O for 3 minutes, 5 times. The sections were placed in filtered thiocarbohydrazide (TCH) solution (10 mg/ml in ddH_2_O) for 20 minutes. The sections were rinsed with ddH_2_O for 3 minutes, 5 times. The sections were treated with 2% osmium oxide for 30 minutes and then washed with ddH_2_O for 3 minutes, 5 times. The sections were placed in 1% (w/v) uranyl acetate at 4°C overnight. The tissue was washed with ddH_2_O for 3 minutes, 5 times and then placed in lead aspartate solution (6.6 mg/ml, pH 5.5) at 60°C for 30 minutes. The tissues were washed in ddH_2_O for 3 minutes, 5 times and then dehydrated in increasing concentration of ethanol (70%, 90%, 95%, 20 minutes once each and then 100% for 10 minutes, 3 times.). Epon embedding was performed with Embed-812 (Electron Microscopy Sciences, 14120), Araldite 502 (Electron Microscopy Sciences, 13900), DDSA (Electron Microscopy Sciences, 13700), DMP-30 (Electron Microscopy Sciences, 13600) in Chien molds and the tissues were left to polymerase overnight at 60°C overnight. The electron micrographs were obtained using Philips EM 410 transmission electron microscope. Fiji was used for electron micrograph image analysis. For synaptic vesicles, only clearly visible vesicles were counted and their areas were measured using freehand selection tool. Post-synaptic density length was measured with freehand line tool. Asymmetric spines and mitochondria were counted per 21000X magnification image; only mitochondria that were visible as a whole were counted, i.e. those that appeared cut along the edges/borders of the image were excluded.

### Glutamate Dehydrogenase Activity Assay

300 µg protein of synaptosomes prepared in Krebs-like buffer were used. Final concentrations of glutamate (Sigma, G1251) and NADP (Sigma, N5755) were 1 mM each. Total final volume per well in the black clear-bottom 96-well plate (Greiner Bio-One, 655090) was 200 µl. Immediately after addition of NADP, the readings were collected at 37°C. Cytation 5 plate reader detected fluorescence every 45 seconds for 1 hour, with excitation and emission wavelengths of 340 and 460 nm, respectively. The data were analyzed using Gen5 software. The slope (rate of product formation) was determined from the linear phase of the fluorescence curve. For the standard curve, various concentrations of glutamate dehydrogenase (Sigma, G2626) were used.

### Glutaminase Activity Assay

5 µg protein of synaptosomes prepared in Krebs-like buffer were used. The assay was carried out according to the manufacturer’s manual (BioVision, K455).

### Aspartate Aminotransferase Activity Assay

5 µg protein of synaptosomes prepared in Krebs-like buffer were used. The assay was carried out according to the manufacturer’s manual (Sigma, MAK455).

### ATP Assay in Synaptosomes

In addition to LC-MS/MS, ATP levels in wild-type and *Gpt2-*null synaptosomes at P18 were quantified with ATP Assay Kit (abcam, 83355, lot: GR3299138) according to the manufacturer’s instructions. The samples were deproteinized by PCA/KOH method as recommended by the manual. Protein amount in each sample was determined using the BCA assay (ThermoScientific Pierce, PI23227).

### Experimental Design and Statistical Analyses

All data in the Figures and the Results section are presented as average ± standard error of the mean, unless otherwise noted. The statistical test for comparisons of two groups was unpaired two-tailed Student t-test unless otherwise noted. All statistical analyses were compiled using GraphPad Prism software (RRID:SCR_002798).

## Results

### GPT2 is the primary glutamate pyruvate transaminase in isolated synaptic terminals

To understand the contribution of GPT2 to glutamate metabolism at the synapse, we conducted studies in slice electrophysiology in parallel with biochemical studies in synaptosomes. Synaptosomes are isolated synaptic terminals that form during homogenization and medium-speed centrifugation of mouse nervous tissues in a sucrose-based solution (Fig. 1A) [20]. GPT2 was enriched both in synaptosomes and mitochondria (Fig. 1C). GPT1, the cytosolic paralog of GPT2, was enriched in the cytosolic fraction but virtually absent in both synaptosomal and mitochondrial fractions. In western blotting of the different subcellular fractions, we also confirmed the presence of some of the proteins involved in glutamate and glutamine metabolism in synaptosomes (Fig. 1B&C). We fractionated synaptosomes further into synaptic membranes, cytosol and vesicles, and GPT2 was not enriched in any of these fractions indicating that GPT2 is enriched in the mitochondria situated in synaptosomes (Fig. 1D). Methods and characterization of synaptosome preparations have been previously detailed [19, 20, 22]. Importantly, while synaptosomes are enriched in pre- and post-synaptic neuronal elements [such as synapsin I (SYNI), synaptic vesicle protein 2 (SV2), vesicular GABA transporter (VGAT), vesicular glutamate transporter (VGLUT1), glutamate ionotropic receptor AMPA Type Subunit 1 (GRIA1)], these isolated synaptic preparations also include astrocyte remnants found at synaptic terminals, as has been noted previously [22–25]. Astrocytic components are reflected by presence of glial fibrillary acidic protein (GFAP) and astrocytic glutamate transporter (GLT1); however, exceedingly low levels of GFAP (Fig. 1C) suggest that astrocytic cytosolic components are quite low, while GLT1 levels may be constituted by astrocytic synaptic membranes as well as neuronal GLT1 [25] (Fig. 1D). Therefore, synaptosomes are used here as an experimental model for isolated synapses; however, this is not meant to indicate that these synaptic preparations are completely devoid of astrocytic components. The model based on our data and the substantial prior literature is illustrated in Fig. 1B.

In agreement with the enrichment of GPT2 as compared to GPT1 in synaptosomes, we found only residual GPT enzyme activity in *Gpt2-*null synaptosomes compared to the wild-type controls (Fig. 1E). To further confirm absence of GPT activity in *Gpt2-*null synaptosomes and assess the function of metabolic pathways that utilize GPT2, we conducted isotope labeling experiments with heavy isotope labeled tracers (Fig. 1F). From amine nitrogen-labeled glutamine ([α-^15^N]-glutamine), alanine was not labeled at all in *Gpt2-*null synaptosome pellets or medium indicating that glutamine could not act as a precursor for alanine in the absence of GPT2. Interestingly, the alanine pool that was released from wild-type synaptosomes into the medium originated from [α-^15^N]-glutamine more than the alanine pool that remained in the pellet (Fig. 1F, left). Akin to a GPT enzyme activity assay, an excess supply of heavy nitrogen labeled alanine (1.6 mM) and unlabeled alpha-ketoglutarate (1.6 mM) resulted in 41.7 ± 2.4% labeling in glutamate of wild-type synaptosomal pellets as well as in 47.2 ± 2.3% of the glutamate pool released into the medium, suggesting that alanine along with alpha-ketoglutarate can be used as substrates for glutamate in our wild-type synaptosome preparations (Fig. 1F, right). On the other hand, *Gpt2-*null synaptosomal pellet or medium showed very low heavy nitrogen labeling in glutamate (5.1 ± 0.4 % in pellet; 4.9 ± 0.4 % in medium). Thus, we conclude that GPT2 is the main alanine aminotransferase in synaptosomes, and that GPT2 can produce a significant fraction of glutamate (approximately 40%) in these isolated synaptic preparations.

### Electrophysiological recordings suggest decreased glutamatergic transmission in *Gpt2-*null hippocampus

To assess the role of GPT2 in synaptic transmission and modulation of glutamate, we conducted whole-cell patch-clamp recording experiments in pyramidal neurons of CA1 hippocampal slices (Fig. 2A). We first characterized the electrophysiological properties of the *Gpt2-*null pyramidal neurons. The resting membrane potential of *Gpt2-*null pyramidal neurons was slightly depolarized (Wild-type: −63.8 ± 1.9 mV vs *Gpt2-*null: −59.8 ± 1.0 mV) (Fig. 2B) and the membrane resistance was increased (Wild-type: 122.7 ± 7.3 MOhm vs *Gpt2-*null: 160.5 ± 9.5 MOhm) (Fig. 2C). The capacitance was reduced (Wild-type: 121.7 ± 6.7 pF vs *Gpt2-*null: 92.7 ± 4.8 pF) (Fig. 2D), suggesting that *Gpt2-*null CA1 pyramidal neurons are smaller in soma size, consistent with our prior studies of *Gpt2*-null neurons [17]. Slight depolarization and increased membrane resistance in *Gpt2-*null pyramidal neurons are visible in the representative traces in Fig. 2A. Overall, the electrophysiological properties of *Gpt2-*null neurons were significantly altered.

**Figure 2.**
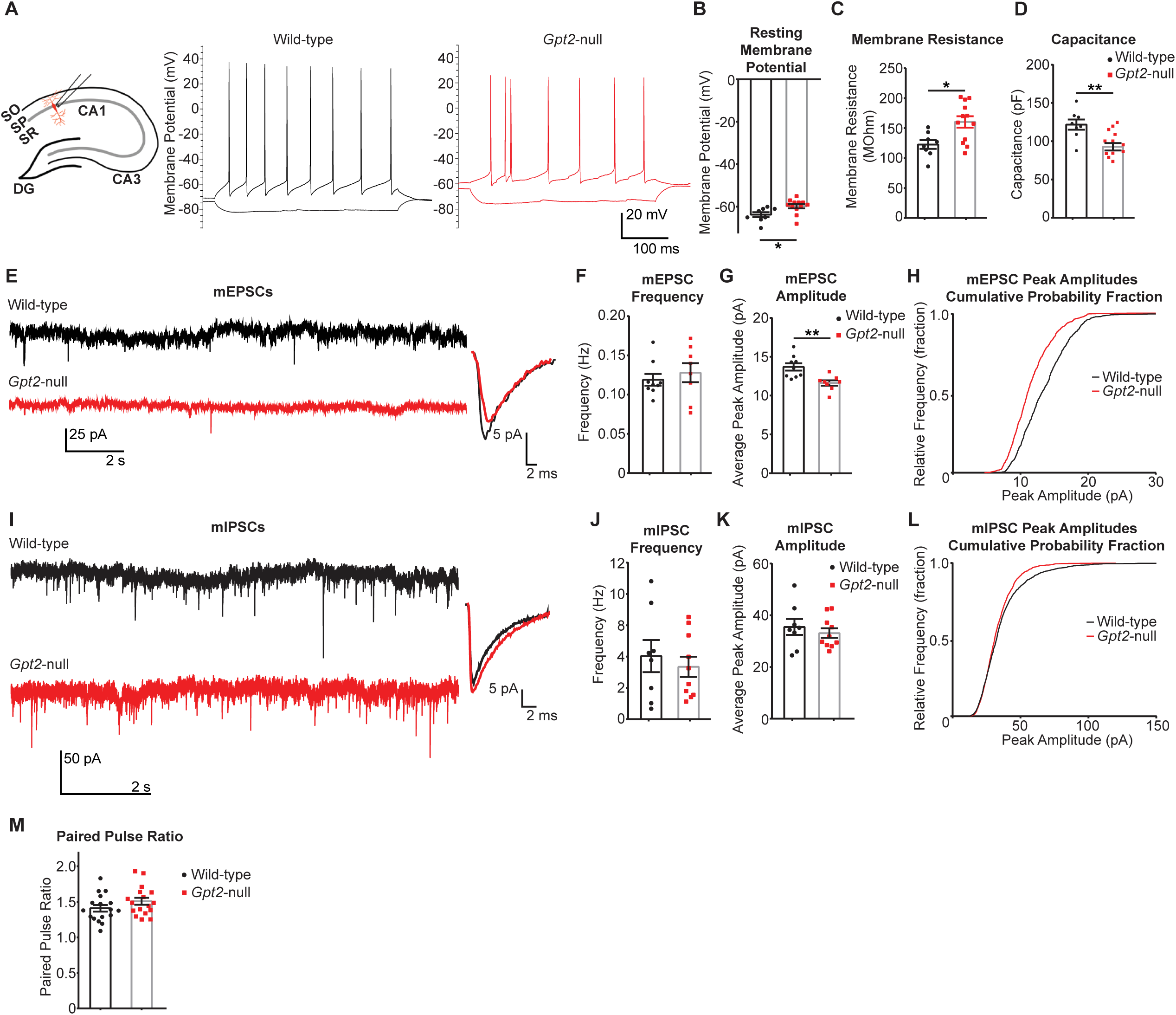
Glutamatergic transmission is diminished in *Gpt2-*null CA1 hippocampus pyramidal neurons. **A.** Schematic of the hippocampal slice preparation and a CA1 pyramidal neuron (red) and recording pipette. The representative whole-cell recordings of wild-type (left, black) and *Gpt2-*null (right, red) CA1 pyramidal neurons in current clamp mode at postnatal day 18 (P18). Current steps of 20 pA were used to depolarize or hyperpolarize the neuron. **B.** Resting membrane potentials of *Gpt2-*null CA1 pyramidal neurons are slightly depolarized. The resting membrane potential of CA1 neurons from wild-type (black) and *Gpt2-*null (red) mice at P18 was obtained immediately after breaking into the cell in current clamp mode. n = 4-5 mice/4-5 slices/8-12 cells. \**P* = 0.02. **C.** *Gpt2-*null CA1 pyramidal neurons have greater membrane input resistance. Each dot represents a different cell recording from wild-type (black) and *Gpt2-*null (red) mice at P18. n = 4-5 mice/4-5 slices/8-12 cells. \**P* = 0.01. **D.** *Gpt2-*null CA1 pyramidal neurons have lower capacitance. Each dot represents a different cell recording from wild-type (black) and *Gpt2-*null (red) mice at P18. \*\**P* = 0.002. **E.** Representative traces of miniature excitatory synaptic currents (mEPSC) from whole-cell recordings in CA1 pyramidal neurons of wild-type (top, black) and *Gpt2-*null (bottom, red) mice at P18. Averaged mEPSCs from one representative cell from wild-type (black) and *Gpt2-*null (red) cells at P18 are shown on the right. **F.** mEPSC frequency is not changed in *Gpt2-*null CA1 pyramidal neurons. Each dot represents a different cell recording from wild-type (black) and *Gpt2-*null (red) mice at P18. n = 4-6 mice/4-6 slices/8-9 cells. *P* = 0.52. **G.** mEPSC peak amplitude is reduced in *Gpt2-*null CA1 pyramidal neurons. Each dot represents a different cell recording from wild-type (black) and *Gpt2-*null (red) mice at P18. n = 4-5 mice/4-5 slices/8-12 cells. \*\**P* = 0.004. **H.** Cumulative probability histogram of mEPSC peak amplitudes from representative wild-type (black) and *Gpt2-*null (red) CA1 pyramidal neurons at P18. Kolmogorov-Smirnov test was conducted, 4 neurons from each genotype were used to construct the curves; total mEPSC events, wild-type: 799 and *Gpt2-*null: 793. \*\*\**P* <0.0001; d = 0.24. **I.** Representative traces of miniature inhibitory synaptic currents (mIPSC) from whole-cell recordings in CA1 pyramidal neurons of wild-type (top, black) and *Gpt2-*null (bottom, red) mice at P18. Averaged mIPSCs from one representative cell from wild-type (black) and *Gpt2-*null (red) CA1 pyramidal neurons at P18 are shown on the right. **J.** mIPSC frequency is not changed in *Gpt2-*null CA1 pyramidal neurons. Each dot represents a different cell recording from wild-type (black) and *Gpt2-*null (red) mice at P18. n = 4-6 mice/4-6 slices/8-9 cells. *P* = 0.56. **K.** mIPSC peak amplitude is not changed in *Gpt2-*null CA1 pyramidal neurons. Each dot represents a different cell recording from wild-type (black) and *Gpt2-*null (red) mice at P18. n = 4-6 mice/4-6 slices/8-9 cells. *P* = 0.5. **L.** Cumulative probability histogram of mIPSC peak amplitudes from representative wild-type (black) and *Gpt2-*null (red) CA1 pyramidal neurons at P18. Kolmogorov-Smirnov test was conducted, 4 neurons from each genotype were used to construct the curves; total mEPSC events, wild-type: 800 and *Gpt2-*null: 800. *P* = 0.1960; D = 0.054. **M.** Paired pulse ratios were unchanged as recorded in *Gpt2*-null CA1 pyramidal neurons at P18. Each dot represents a different cell recording from wild-type (black) and *Gpt2*-null (red) mice at P18. n = 7-9 mice/17 slices/17 cells. *P* = 0.16.

Miniature post-synaptic currents are indirect measures of neurotransmitter levels in synaptic quanta, individual synaptic vesicles [26, 27]. We recorded miniature post-synaptic excitatory currents (mEPSCs) from *Gpt2-*null CA1 pyramidal neurons voltage-clamped at −80mV. The mEPSC frequency was unchanged (Fig. 2F, Wild-type: 0.12 ± 0.01 Hz vs *Gpt2-*null: 0.13 ± 0.01 Hz); however, the mEPSC peak amplitude was decreased in *Gpt2-*null pyramidal neurons (Fig. 2E&G, Wild-type: 13.7 ± 0.5 pA vs *Gpt2-*null: 11.6 ± 0.3 pA). The cumulative probability histogram of mEPSC peak amplitudes of all miniature excitatory events was shifted to the left in a representative *Gpt2-*null pyramidal neuron compared to its wild-type control (Fig. 2H, Kolmogorov-Smirnov D value: 0.2386). Overall, these results are consistent with a model that glutamate levels are reduced in synaptic vesicles of excitatory afferents on *Gpt2-*null CA1 pyramidal neurons.

To test whether the GABAergic input onto GPT2 pyramidal neurons was similarly affected, we recorded miniature post-synaptic inhibitory currents (mIPSCs) from *Gpt2-* null CA1 pyramidal neurons voltage-clamped at −80mV (Fig. 2I-K). Both mIPSC frequency (Fig. 2J) and peak amplitude (Fig. 2K) were unchanged in *Gpt2-*null CA1 pyramidal neurons (Wild-type: 4.04 ± 1.03 Hz vs *Gpt2-*null: 3.3 ± 0.65 Hz; Wild-type: 35.5 ± 3.0 pA vs *Gpt2-*null: 33.1 ± 1.9 pA, respectively). The cumulative probability histogram of mIPSC peak amplitudes had a similar distribution in the wild-type control (Fig. 2L). Together, these results suggest that GABA levels remain the same in synaptic vesicles at inhibitory synapses on *Gpt2-*null CA1 pyramidal neurons.

To rule out any changes in synaptic release probability, we examined paired pulse ratios in CA1 pyramidal neurons. The paired pulse facilitation remained the same in *Gpt2-*null pyramidal neurons (Figure 2M, Wild-type: 1.41 ± 0.05 vs *Gpt2-*null: 1.51 ± 0.05) suggesting that the synaptic release probability is unchanged.

### Released glutamate levels are reduced in *Gpt2-*null synaptosomes

To determine the extent to which vesicular glutamate levels were decreased in *Gpt2-*null synaptosomes and to corroborate the electrophysiology data, we measured glutamate release upon depolarization by liquid chromatography/mass spectrometry (LC/MS). We initially confirmed the viability of our synaptosome preparations by observing increased levels of glutamate released after depolarization with 50 mM KCl in Krebs-like buffer compared to the release at baseline (Fig. 3A). Overall glutamate levels in the cytosolic fraction of the *Gpt2-*null forebrain obtained during synaptosome preparation were significantly reduced as determined by enzymatic detection (Fig. 3B). Overall glutamate levels in total *Gpt2*-null synaptosome preparations were also reduced compared to their wild-type controls (Fig. 3C). We then tested glutamate release and discovered that levels of glutamate released upon KCl stimulation were reduced in *Gpt2-*null synaptosomes (Fig. 3D). In contrast, released GABA levels were unchanged (Fig. 3E) which agrees with the unchanged mIPSC amplitudes in CA1 hippocampal slices (Fig. 2K). We also tested for changes in protein levels of vesicular glutamate transporter (VGLUT1) and vesicular GABA transporter (VGAT) in *Gpt2*-null synaptosomes (Fig. S1). Interestingly, we see increases in both VGLUT1 and VGAT in *Gpt2*-null synaptosomes suggesting that decreases of glutamate release are unlikely to be due to changes in protein levels of vesicular transporters.

**Figure 3.**
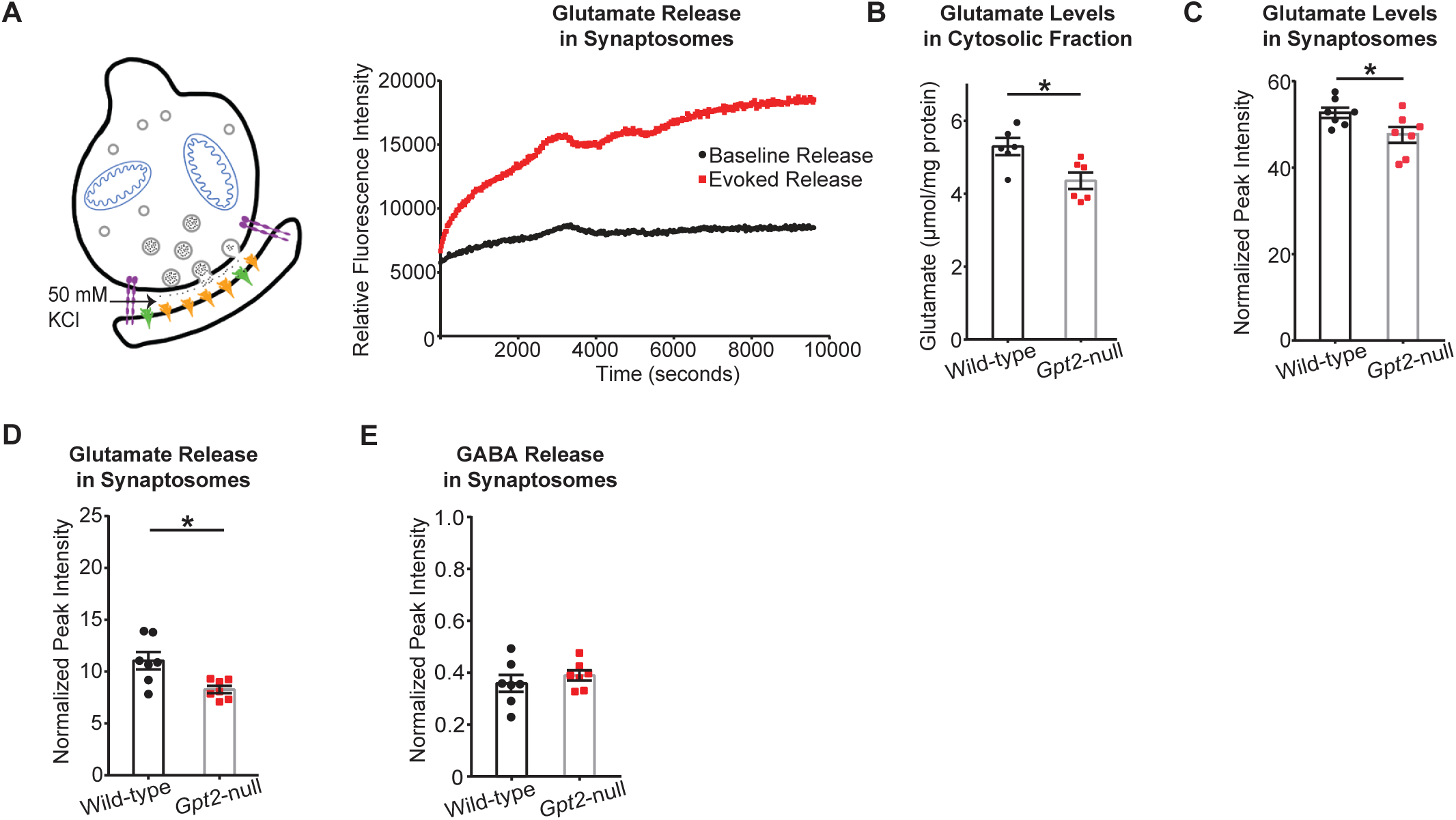
Glutamate released upon depolarization is reduced in *Gpt2-*null synaptosomes. **A.** Schematic of the synaptosomes and evoking glutamate release from synaptic vesicles with 50 mM KCl. A representative trace of relative fluorescence intensity across time is shown (collected at excitation and emission wavelengths of 340 and 460 nm, respectively). Viability of synaptosomes was confirmed by adding 50 mM KCl to synaptosomes and comparing baseline (black) and evoked (red) glutamate levels in wild-type synaptosomes obtained at postnatal day 18 (P18). **B.** Overall glutamate levels in the cytosolic fraction are reduced in *Gpt2-*null mice. Cytosol fraction of the forebrain was obtained by separating the supernatant from the crude synaptosomal pellet. Glutamate was detected by enzymatic method and expressed as µmol per mg of protein. Each dot represents a different synaptosome sample obtained from wild-type (black) or *Gpt2-*null (red) mice at P18. \**P* = 0.02. **C.** Overall glutamate levels are reduced in *Gpt2*-null synaptosomes. Each dot represents a different synaptosome sample obtained from wild-type (black) or *Gpt2-*null (red) mice at P18. \**P* = 0.037. **D.** Glutamate levels released upon depolarization are reduced in *Gpt2-*null synaptosomes. Each dot represents a different synaptosome sample obtained from wild-type (black) or *Gpt2-*null (red) mice at P18. \**P* = 0.01. **E.** Released GABA levels were unchanged in *Gpt2-*null synaptosomes. Each dot represents a different synaptosome sample obtained from wild-type (black) or *Gpt2-*null (red) mice at P18. *P* = 0.44.

Glutamate metabolism is influenced strongly by glutaminase [3], which converts glutamine into glutamate in the mitochondria and by the most abundant aminotransferase in the brain, aspartate aminotransferase [28], which catalyzes a reversible transamination to interconvert aspartate to glutamate. Given this influence on glutamate, we checked whether any alterations in enzyme activity or protein levels of these enzymes may contribute to the decreases in glutamate. Aspartate levels in *Gpt2*-null synaptosomes were unchanged (Fig. S2A) as well as aspartate aminotransferase enzyme activity (Fig. S2B). There were no changes in ^15^N labeling in aspartate using [α-^15^N]-glutamine as tracer (Fig. S2C), however, expectedly, zero ^15^N labeling in aspartate was observed when [α-^15^N]-alanine was used as tracer as this route would be unavailable due to the loss of GPT2 (Fig. S2C). Similarly, glutaminase enzyme activity in *Gpt2*-null synaptosomes was unaltered (Fig. S2D). Protein levels of aspartate aminotransferase and glutaminase in *Gpt2*-null hippocampus tissue were similar to those of their wild-type controls (Fig. S2E). Overall, the data suggests that alterations of glutamate in *Gpt2*-null synaptosomes were not primarily caused by changes in either aspartate aminotransferase or glutaminase activity.

### Increased synaptic vesicle size in *Gpt2-*null CA1 stratum radiatum

To investigate whether ultrastructural changes may be associated with the decreased glutamatergic transmission (Fig. 2), we examined glutamatergic synapses based on the identification of asymmetric spines in *Gpt2-*null CA1 stratum radiatum by electron microscopy (Fig. 4 and S3). Interestingly, we observed approximately 10% increase in area of individual synaptic vesicles in *Gpt2-*null CA1 asymmetric spine synapses (Wild-type: 1361 ± 3.1 nm^2^ vs. *Gpt2-*null: 1535 ± 8.7 nm^2^, *P*=0.0013) (Fig. 4B). Number of synaptic vesicles per synapse (Fig. 4C) and post-synaptic density length (Fig. 4D) were similar to the wild-type controls. Spine counts as well as mitochondria counts per image were unchanged in *Gpt2-*null CA1 stratum radiatum (Fig. S3B&C). Overall, the electron micrographs indicate that the ultrastructural morphology in *Gpt2-*null CA1 stratum radiatum was intact, albeit with a significant increase in synaptic vesicle size.

**Figure 4.**
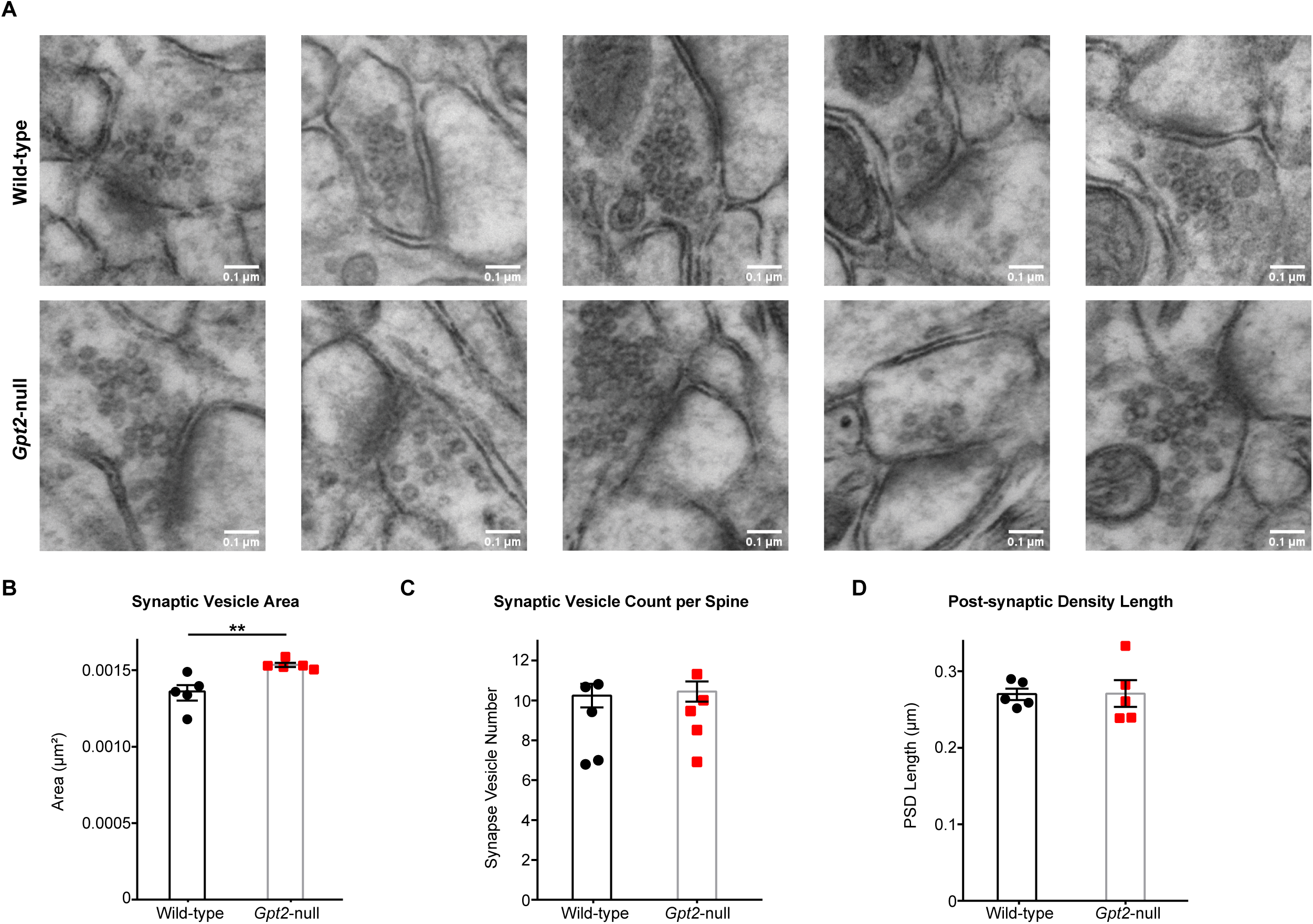
Synaptic vesicle area is increased in *Gpt2-*null CA1 stratum radiatum while number of synaptic vesicles per spine remains unchanged. **A.** Representative images of spines in wild-type (top) and *Gpt2-*null (bottom) CA1 stratum radiatum at P18. Scale bar: 0.1 µm. **B.** Quantification of the area of synaptic vesicles. Each dot represents the averaged area of synaptic vesicles in spines obtained from one animal. A minimum of 25 micrographs were used for each animal. \*\**P* = 0.001. **C.** Quantification of the number of synaptic vesicles per spine. Each dot represents the averaged number of synaptic vesicles obtained from one animal. A minimum of 25 micrographs were used for each animal. **D.** Quantification of the post-synaptic density length. Each dot represents the averaged post-synaptic density length obtained from one animal. A minimum of 25 micrographs were used for each animal.

### Amelioration of glutamate release by alpha-ketoglutarate supplementation in *Gpt2-*null synaptosomes

We hypothesized that synaptic glutamate release may be ameliorated by alpha-ketoglutarate. This hypothesis is based in part on prior observations that TCA cycle intermediates, particularly, alpha-ketoglutarate have been shown to be precursors for synaptic glutamate via several metabolic reactions [29–32]. Therefore, we sought to rescue the decreases in synaptic glutamate levels seen in *Gpt2-*null synaptosomes by supplementing with alpha-ketoglutarate. When supplemented with alpha-ketoglutarate, released glutamate levels of *Gpt2-*null synaptosomes were restored to the wild-type levels (Fig. 5A). Alanine, another major product of the GPT2 reaction, failed to correct the synaptic glutamate levels (Fig. 5B). Alpha-ketoglutarate combined with alanine also corrected the glutamate levels in *Gpt2-*null synaptosomes; however, to near equivalent levels to alpha-ketoglutarate alone (Fig. 5C). In contrast, released GABA levels in *Gpt2-*null synaptosomes were unchanged as compared to the wild-type, with or without alpha-ketoglutarate or alanine supplementation (Fig. 5D-F). As a control, we also demonstrated that alanine and alpha-ketoglutarate readily enter synaptosomes (Fig. S4A-D). Notably, baseline alpha-ketoglutarate levels were unchanged but alanine levels were reduced in *Gpt2-*null synaptosomes (Fig. S4A-D) in corroboration of our previous finding that alanine levels in the whole *Gpt2-*null brain were reduced [11, 33]. Overall, these results indicate that alpha-ketoglutarate was able to rescue glutamate levels and that a metabolic intervention was sufficient to correct glutamate deficiency in *Gpt2-*null synaptosomes.

**Figure 5.**
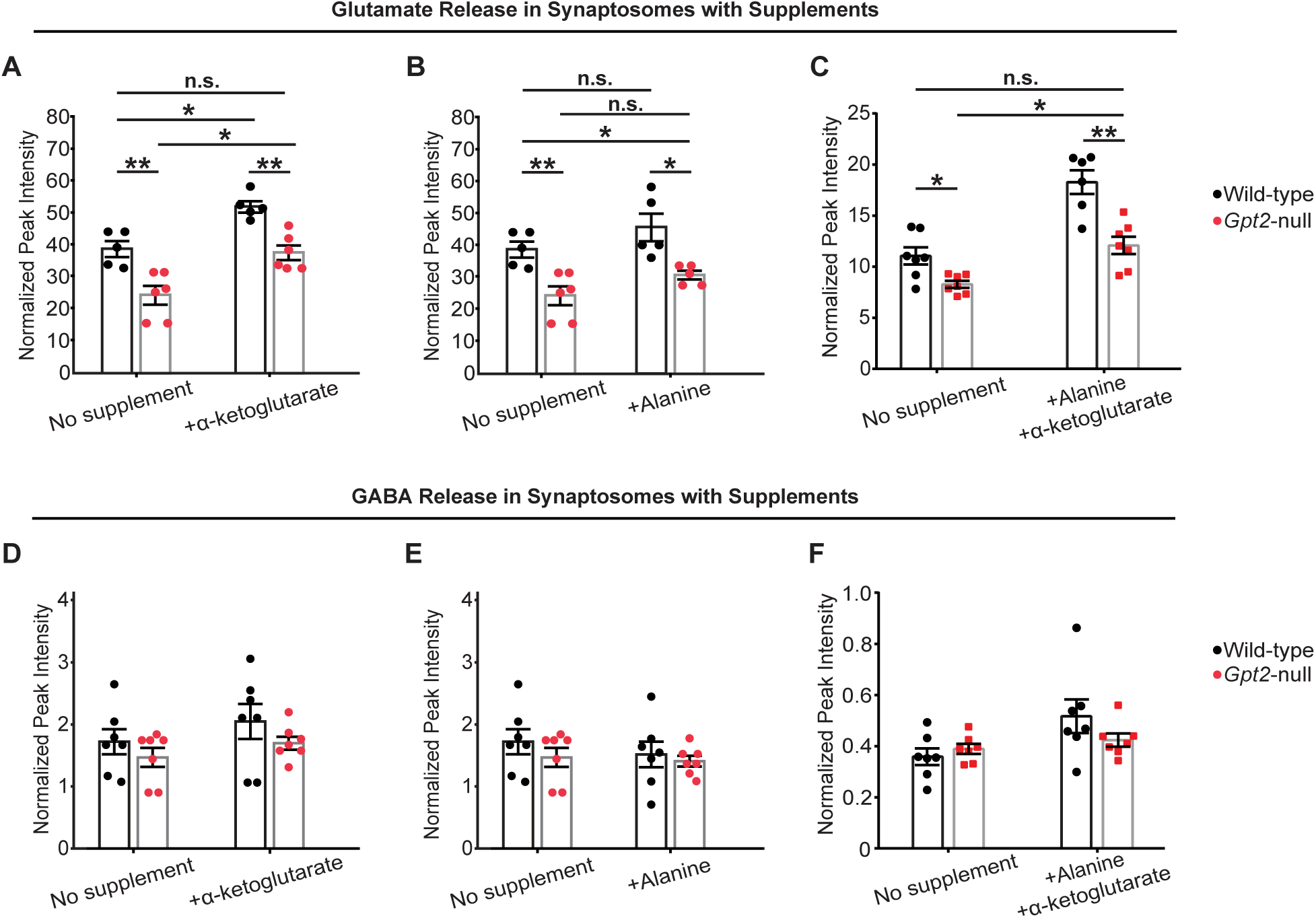
Decreases in released glutamate are restored by alpha-ketoglutarate in *Gpt2-*null synaptosomes. **A.** Glutamate levels are restored by alpha-ketoglutarate in *Gpt2-*null synaptosomes. Each dot represents a different synaptosome sample obtained from wild-type (black) or *Gpt2-*null (red) mice at P18. Wild-type vs. *Gpt2-*null no supplement: \*\*\**P* < 0.001; +alpha-ketoglutarate: \*\*\**P* < 0.001. Wild-type no supplement vs. *Gpt2-*null +alpha-ketoglutarate: *P* = 0.99. Wild-type no supplement vs. wild-type +alpha-ketoglutarate: \*\*\**P* < 0.001. Wild-type vs. *Gpt2-*null +alpha-ketoglutarate: \*\*\**P* < 0.001. n.s. = statistically non-significant. A two-way ANOVA was performed with Tukey’s multiple comparison correction and the reported p-values are adjusted p-values. **B.** Glutamate levels are not restored by alanine in *Gpt2-*null synaptosomes. Each dot represents a different synaptosome sample obtained from wild-type (black) or *Gpt2-*null (red) mice at P18. Wild-type vs. *Gpt2-*null no supplement: \*\*\**P* < 0.001; +alanine: \**P* = 0.01. Wild-type no supplement vs. *Gpt2-*null +alanine: \**P* = 0.02. Wild-type no supplement vs. alanine: *P* = 0.16. *Gpt2-*null no supplement vs. alanine: P = 0.10. n.s. = statistically non-significant. A two-way ANOVA was performed with Tukey’s multiple comparison correction and the reported p-values are adjusted p-values. **C.** Glutamate levels are restored by alanine and alpha-ketoglutarate combined supplementation in *Gpt2-*null synaptosomes. Each dot represents a different synaptosome sample obtained from wild-type (black) or *Gpt2-*null (red) mice at P18. Wild-type vs. *Gpt2-*null no supplement: \*\*\**P* < 0.001. Wild-type vs. *Gpt2-*null +alanine +alpha-ketoglutarate: \*\*\**P* < 0.001. Wildtype no supplement vs. *Gpt2-*null +alanine +alpha-ketoglutarate: *P* = 0.84. Wild-type no supplement vs. +alanine+alpha-ketoglutarate: \*\*\**P* < 0.001. *Gpt2-*null no supplement vs. +alanine+alpha-ketoglutarate: \*\*\**P* < 0.001. n.s. = statistically non-significant. A two-way ANOVA was performed with Tukey’s multiple comparison correction and the reported p-values are adjusted p-values. **D.** Released GABA levels were unchanged with or without alpha-ketoglutarate in *Gpt2-* null synaptosomes. Each dot represents a different synaptosome sample obtained from wild-type (black) or *Gpt2-*null (red) mice at P18. Wild-type vs. *Gpt2-*null no supplement: *P* = 0.59. Wild-type vs. *Gpt2-*null +alpha-ketoglutarate: *P* = 0.23. A two-way ANOVA was performed with Tukey’s multiple comparison correction and the reported p-values are adjusted p-values. **E.** Released GABA levels with or without alanine. Each dot represents a different synaptosome sample obtained from wild-type (black) or *Gpt2-*null (red) mice at P18. Wild-type vs. *Gpt2-*null no supplement: *P* = 0.59. Wild-type vs. *Gpt2-*null +alanine *P* = 0.63. A two-way ANOVA was performed with Tukey’s multiple comparison correction and the reported p-values are adjusted p-values. **F.** Released GABA levels with or without alanine and alpha-ketoglutarate combined supplementation. Each dot represents a different synaptosome sample obtained from wild-type (black) or *Gpt2-*null (red) mice at P18. Wild-type vs. *Gpt2-*null no supplement: 0.87. Wild-type vs. *Gpt2-*null +alanine +alpha-ketoglutarate *P* = 0.69. A two-way ANOVA was performed with Tukey’s multiple comparison correction and the reported p-values are adjusted p-values. Data in panel A&B&D&E were derived from the same cohort of animals, and data in panel C&F were derived from a separate cohort of animals.

### Glutamine metabolism is altered in *Gpt2-*null synaptosomes and rescued by alpha-ketoglutarate supplementation

Glutamate is replenished in neurons primarily by the glutamate-glutamine cycle at the tripartite synapse, formed by the pre-synaptic neuron, post-synaptic neuron and astrocyte compartments [34] (Fig. 6A). We observed changes in glutamine metabolism in *Gpt2-*null synaptosomes which we predict are compensatory reflecting loss of GPT2 enzyme activity. First, we observed increased glutamine levels in *Gpt2-*null synaptosomes (Fig. 6B). Second, we discovered an increased fraction of double heavy nitrogen labeled glutamine (Fig. 6C) that is glutamine with a heavy nitrogen in both amine and amide groups. Increased double labeled glutamine suggested upregulation of glutamate metabolizing enzymes, specifically glutamate dehydrogenase and glutamine synthetase. Given the rescue of glutamate levels by alpha-ketoglutarate, we tested if alpha-ketoglutarate would correct this alteration in glutamine metabolism, particularly since alpha-ketoglutarate as a substrate has been shown to suppress glutamate dehydrogenase activity [35]. Indeed, we observed that the fractional enrichment of double-labeled glutamine was reduced back to wild-type levels by alpha-ketoglutarate supplementation in *Gpt2-*null synaptosomes (Fig. 6C). In contrast, alanine failed to reduce double labeling of glutamine in *Gpt2-*null synaptosomes. Glutamine entry into synaptosomes was not affected by alpha-ketoglutarate as evident in unchanged glutamine levels (Fig. S5A) as well as in unchanged fractional enrichments of single heavy nitrogen labeled glutamine (Fig. S5B) and glutamate (Fig. S5C) compared to the no supplement condition. This suggests that the action of alpha-ketoglutarate in decreasing the fractional enrichment of double-labeled glutamine was not caused by decreases in glutamine entry but by changes in glutamine metabolism. With alanine supplementation, overall glutamine levels were decreased approximately by 40% in both wild-type and *Gpt2-*null synaptosomes (Fig. S5A). While glutamine entry may be affected by alanine, as they compete for the same neutral amino acid transporters [36], this led to only modest decreases in the fractional enrichment of single heavy nitrogen labeled glutamine (Fig. S5B) and glutamate (Fig. S5C). This is also reflected by the observation that glutamate levels were unchanged with alanine supplementation (Fig. 5B). Furthermore, alanine supplement did not result in any changes in fractional enrichment of double labeled glutamine in *Gpt2-*null synaptosomes (Fig. 6C). Overall, these results suggest that the reductions in double-labeled glutamine are feasible by alpha-ketoglutarate and not alanine.

**Figure 6.**
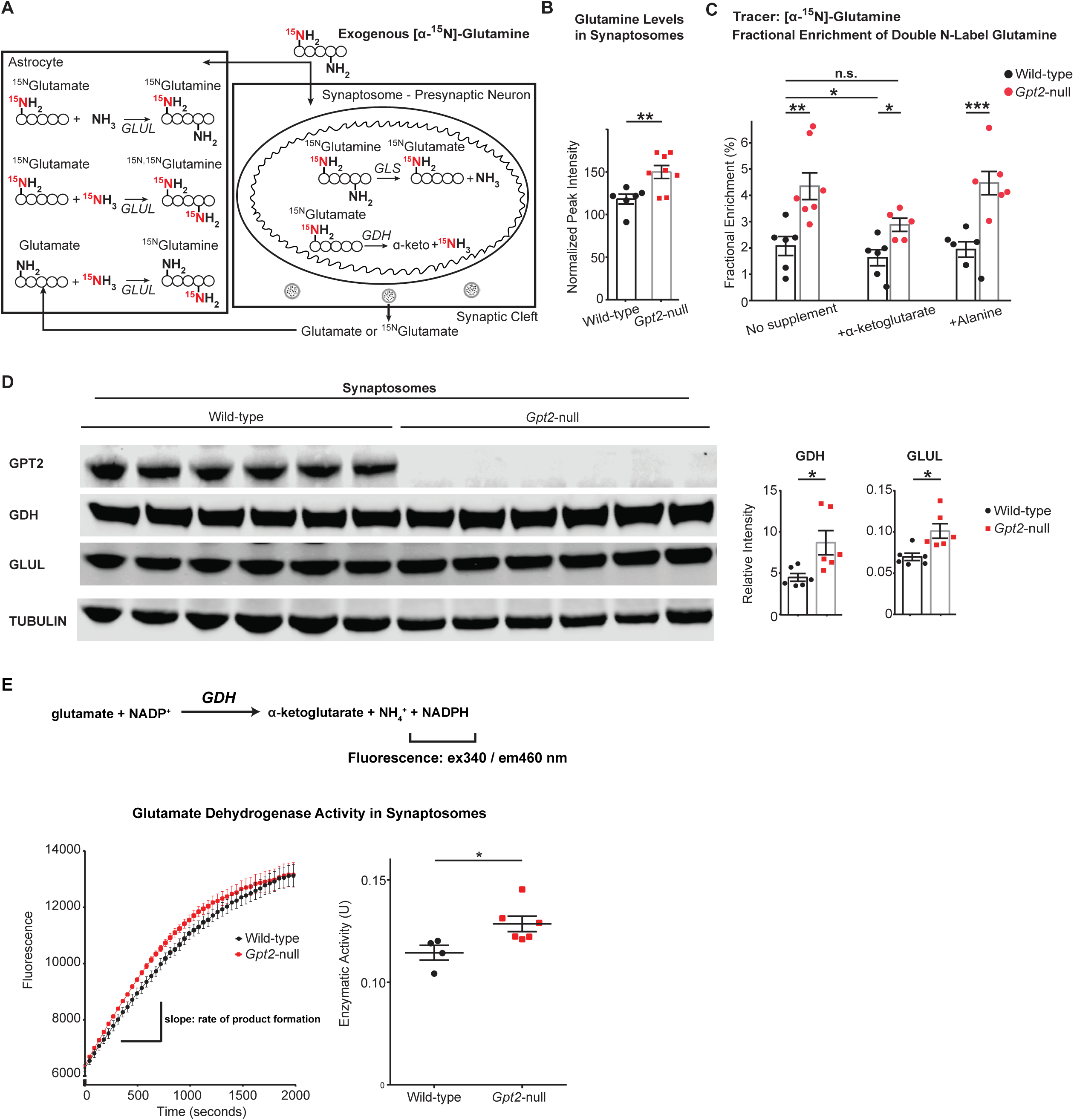
Glutamine metabolism is altered in *Gpt2-*null synaptosomes and rescued by alpha-ketoglutarate. **A.** Metabolic pathways relevant to glutamate/glutamine metabolism and isotope labeling patterns originating from exogenous amine nitrogen labeled glutamine in synaptosomes. *GLS*: glutaminase, *GDH*: glutamate dehydrogenase. *GLUL*: glutamine synthetase. **B.** Baseline glutamine levels are increased in *Gpt2-*null synaptosomes. Each dot represents synaptosome samples obtained from different wild-type (black) or *Gpt2-*null (red) mice at P18. Wild-type vs. *Gpt2-*null \*\**P* = 0.009. **C.** Fractional enrichment of double-labeled glutamine is increased in *Gpt2-*null synaptosomes and restored by alpha-ketoglutarate. Each dot represents a different synaptosome sample obtained from wild-type (black) or *Gpt2-*null (red) mice at P18. Wild-type vs. *Gpt2-*null no supplement: \*\*\**P* < 0.001; +alpha-ketoglutarate: \*\*\**P* < 0.001; +alanine: \*\*\**P* < 0.001; Wild-type no supplement vs. *Gpt2-*null +alpha-ketoglutarate: *P* = 0.31. A two-way ANOVA was performed with Tukey’s multiple comparison correction and the reported p-values are adjusted p-values. **D.** Glutamate dehydrogenase (GDH) and glutamine synthetase (GLUL) protein levels are increased in *Gpt2*-null synaptosomes. Western blotting of wild-type and *Gpt2-*null synaptosome protein lysates collected at P18. Note that GPT2 is absent in *Gpt2-*null (MUT) but present in wild-type (WT) protein lysates. Each dot represents a different protein lysate sample from wild-type (black) or *Gpt2-*null (red) mice. Each protein band is normalized to its corresponding intensity of the actin band. Wild-type vs. *Gpt2-*null: \**P* (GDH, glutamate dehydrogenase) = 0.02; \**P* (GLUL, glutamine synthetase) = 0.01. **E.** Glutamate dehydrogenase enzymatic activity is increased in *Gpt2-*null synaptosomes. Chemical reaction catalyzed by glutamate dehydrogenase (GDH) is shown on top. The enzyme activity assay detects fluorescence with excitation and emission wavelengths of 340 and 460 nm, respectively. *Gpt2-*null synaptosomes display higher GDH activity compared to wild-type controls. The fluorescence of NAPDH (product) over time is shown on the left. The enzymatic activity is determined by the slope of the linear phase, shown on the right. Each dot represents a synaptosomes sample obtained from a different animal. \**P* = 0.03.

Double-labeled glutamine can be formed by the concerted actions of glutamate dehydrogenase and glutamine synthetase (Fig. 6A). We hypothesized that expression and activity of these metabolic enzymes could be upregulated in *Gpt2-*null synaptosomes and that this upregulation can explain the increases in fractional enrichment of double-labeled glutamine in *Gpt2-*null synaptosomes. We performed western blotting in synaptosomes (Fig. 6D) for these metabolic enzymes. We find increases of glutamate dehydrogenase and glutamine synthetase in *Gpt2-*null synaptosomes. These results suggest that increased protein levels of glutamate dehydrogenase and glutamine synthetase may lead to increases in double-labeled glutamine. Complementary to western blotting, we directly tested glutamate dehydrogenase activity by an enzyme activity assay (Fig. 6E, Fig. S6). *Gpt2-*null synaptosomes had significantly more glutamate dehydrogenase activity compared to wild-type controls (Fig. 6E). This finding is in agreement with the increased protein levels of glutamate dehydrogenase and the increased fractional enrichment of double-labeled glutamine in *Gpt2-*null synaptosomes.

### TCA cycle deficits in *Gpt2-*null synaptosomes are rescued by alpha-ketoglutarate supplementation

GPT2 and glutamate dehydrogenase not only modulate glutamate levels but also replenish the tricarboxylic acid (TCA) cycle by augmenting alpha-ketoglutarate [37]. Loss of GPT2 leads to deficits in the TCA cycle intermediates in the mouse brain, likely through impaired anaplerosis [11, 38]. We investigated whether similar TCA cycle deficits were present in *Gpt2-*null synaptosomes (Fig. 7). We observed decreases in baseline levels of malate and fumarate (Fig. 7A). In a separate experiment, we tested whether alpha-ketoglutarate would be able to rescue deficits in malate (Fig. 7B). Malate levels were corrected by alpha-ketoglutarate in *Gpt2-*null synaptosomes.

**Figure 7.**
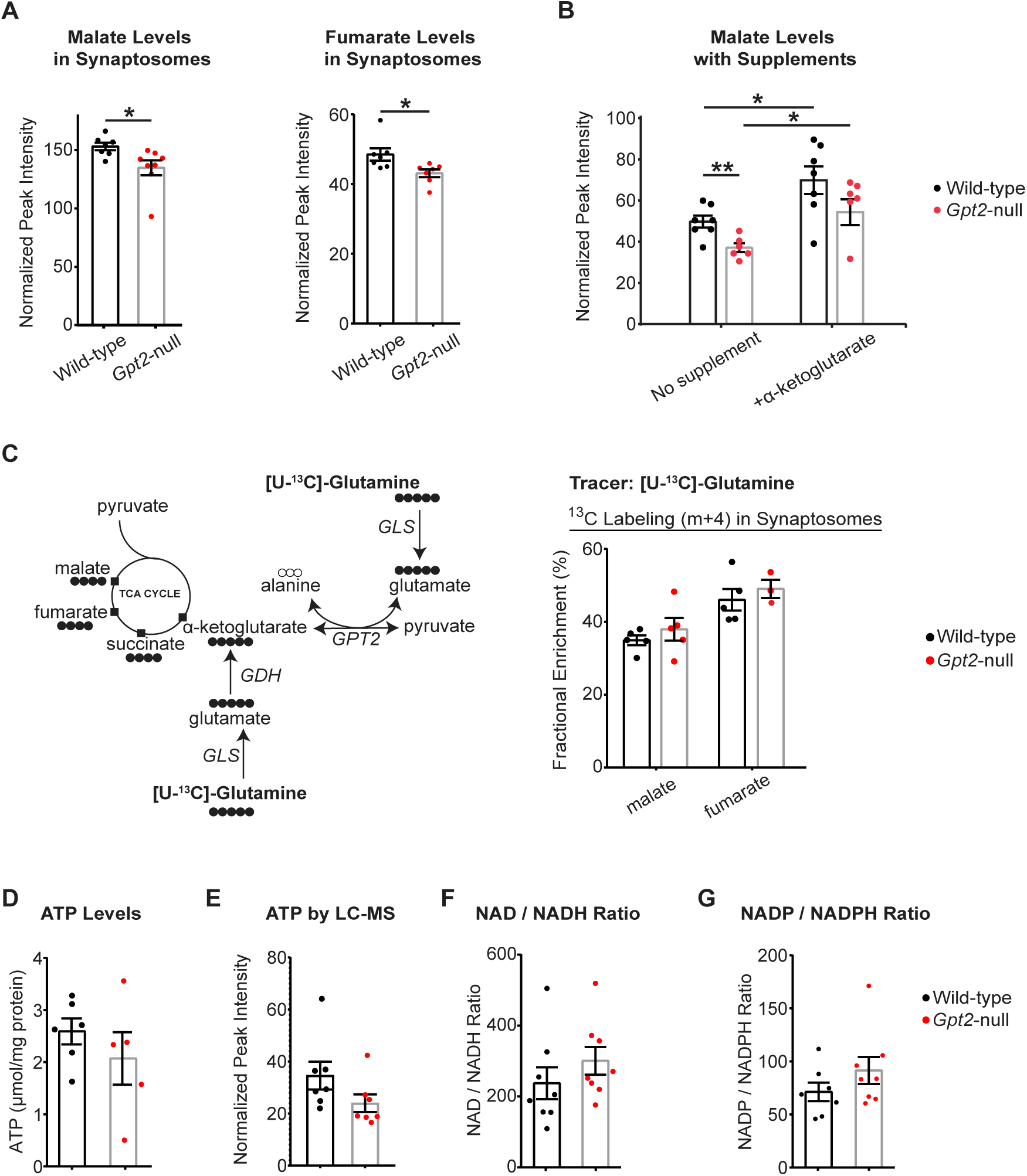
TCA cycle intermediates are reduced but glutamine entry into the TCA cycle is intact in *Gpt2-*null synaptosomes. **A.** Baseline malate and fumarate levels are reduced in *Gpt2-*null synaptosomes. Each dot represents a different sample from wild-type (black) or *Gpt2-*null (red) mice at P18. Baseline wild-type vs. *Gpt2-*null malate: \**P* = 0.03; fumarate: \**P* = 0.03. **B.** Malate levels are rescued by alpha-ketoglutarate in *Gpt2-*null synaptosomes. Each dot represents a different sample from wild-type (black) or *Gpt2-*null (red) mice at P18. Malate wild-type vs. *Gpt2-*null no supplement: \*\**P* = 0.003; wild-type no supplement vs. +alpha-ketoglutarate: \*\**P* = 0.002; *Gpt2-*null no supplement vs. +alpha-ketoglutarate: \*\**P* = 0.002; Wild-type no supplement vs. *Gpt2-*null +alpha-ketoglutarate: *P* = 0.84. A two-way ANOVA was performed with Tukey’s multiple comparison correction and the reported p-values are adjusted p-values. **C.** Fractional enrichments of carbon labeled TCA cycle intermediates using the uniformly carbon labeled [U-^13^C]-Glutamine precursor is unchanged in *Gpt2-*null synaptosomes. The metabolic pathways that label the carbons of TCA cycle intermediates are shown on the left. Fractional enrichments of m+4 (4 heavy carbons) as a percentage are given on the right. Each dot represents synaptosome samples obtained from different wild-type (black) or *Gpt2-*null (red) mice at P18. Wild-type vs. *Gpt2-*null malate: *P* = 0.83; fumarate: *P* = 0.88. A two-way ANOVA was performed with Tukey’s multiple comparison correction and the reported p-values are adjusted p-values. **D.** ATP levels are unchanged in *Gpt2-*null synaptosomes. Each dot represents synaptosome samples obtained from different wild-type (black) or *Gpt2-*null (red) mice at P18. ATP levels were enzymatically determined. Wild-type vs. *Gpt2-*null: *P* = 0.35. **E.** ATP levels as determined by LC-MS. Each dot represents synaptosome samples obtained from different wild-type (black) or *Gpt2-*null (red) mice at P18. ATP levels were enzymatically determined. Wild-type vs. *Gpt2-*null: *P* = 0.12. **F.** NAD to NADH ratio is unchanged in *Gpt2-*null synaptosomes. Each dot represents normalized peak intensities of synaptosome samples obtained from different wild-type (black) or *Gpt2-*null (red) mice at P18 as determined by LC-MS. Wild-type vs. *Gpt2-*null: *P* = 0.31. **G.** NADP to NADPH ratio is unchanged in *Gpt2-*null synaptosomes. Each dot represents normalized peak intensities of synaptosome samples obtained from different wild-type (black) or *Gpt2-*null (red) mice at P18 as determined by LC-MS. Wild-type vs. *Gpt2-*null: *P* = 0.23.

We also tested for glutamine as a precursor for the TCA cycle intermediates in *Gpt2-*null synaptosomes (Fig. 7C). Given the loss of GPT2, a decreased fractional enrichment of labeled TCA cycle intermediates using [U-^13^C]-glutamine as precursor was expected (Fig. 7C, left). However, the fractional enrichments for the TCA cycle intermediate measured were unchanged in *Gpt2-*null synaptosomes (Fig. 7C, right). These results indicate that glutamine may be used by *Gpt2-*null synaptosomes to supplement the TCA cycle. Finally, we also tested for other metrics of energetics such as ATP levels as well as NAD to NADH and NADP to NADPH ratios (Fig. 7D-G). All remained unchanged in *Gpt2-*null synaptosomes. Overall, these results on TCA cycle suggest that the TCA intermediates are decreased; however, energetics may be approximately maintained. This compensation may be due to upregulation of metabolic enzymes (Fig. 6), such as glutamate dehydrogenase, which may provide an alternative carbon source to the TCA cycle in *Gpt2-*null synaptosomes.

## Discussion

Glutamate metabolism is maintained and tightly regulated by a concert of metabolic enzymes and substrate transporters [37, 39–42]. An imbalance of substrates or proteins involved in glutamate metabolism often leads to a compensatory mechanism that takes advantage of alternative metabolic pathways [43–48]. This flexibility of the metabolome related to glutamate is realized by 44 distinct enzymes that consume or produce glutamate, as curated in Kyoto Encyclopedia of Genes and Genomes (KEGG) [49]. Here, we report that loss of glutamate pyruvate transaminase 2 (GPT2) may cause impaired glutamatergic transmission and compensatory changes in proteins related to glutamate and glutamine metabolism. The observed defect in glutamate release from synaptosomes and other metabolic defects, such as in glutamine and TCA metabolism, were alleviated by alpha-ketoglutarate supplementation. While the glutamate-glutamine cycle has garnered substantial study as mechanisms that ensures glutamate availability for excitatory neurotransmission, our study of *Gpt2-*null mice and a growing literature draw attention to the interplay between pre-synaptic mitochondrial TCA cycle metabolism and glutamatergic synaptic transmission [10]. In addition, our study has potential relevance to the origin of cognitive disability in children with GPT2 Deficiency.

We demonstrate that GPT2 is the primary glutamate pyruvate transaminase in synaptosomes, and that GPT2 is localized to mitochondria and enriched in synaptosomes (Fig. 8). GPT2 supports cellular growth by allowing glutamate oxidation in mitochondria thereby enhancing biosynthesis and energetics (Fig. 8A) [11, 33, 50, 51]. In agreement with this, GPT2 deficiency results in smaller soma sizes and slightly depolarized CA1 pyramidal neurons (Fig. 2) as well as microcephaly in both mice and human patients with *GPT2* mutations as previously demonstrated [11]. Glutamate, the major excitatory neurotransmitter, is an excellent source of carbon for the TCA cycle in mitochondria [3, 4, 52, 53]. We find deficits in the TCA cycle of *Gpt2-*null synaptosomes including decreases in malate and fumarate levels (Fig. 8B, event 2). Loss of GPT2 may lead to upregulation of other metabolic enzymes that allow for intact glutamine entry into the TCA cycle in the absence of GPT2. In *Gpt2-*null synaptosomes, we observe increased protein and activity levels of glutamate dehydrogenase (GDH), which deaminates glutamate forming alpha-ketoglutarate and ammonia (Fig. 8B, event 3), along with increases in glutamine synthetase (GLUL) protein levels which allows for efficient ammonia fixation. This may explain the increases in fractional enrichment of double heavy nitrogen labeled glutamine and in overall glutamine levels. GDH catalyzes practically an irreversible reaction that favors glutamate deamination, and thus once glutamate is consumed by this enzyme, it is fixed to its keto acid, alpha-ketoglutarate [47, 54, 55]. Elevations of glutamate dehydrogenase activity may provide an explanation as to why glutamate levels tend to decrease in *Gpt2-*null brain tissue.

**Figure 8.**
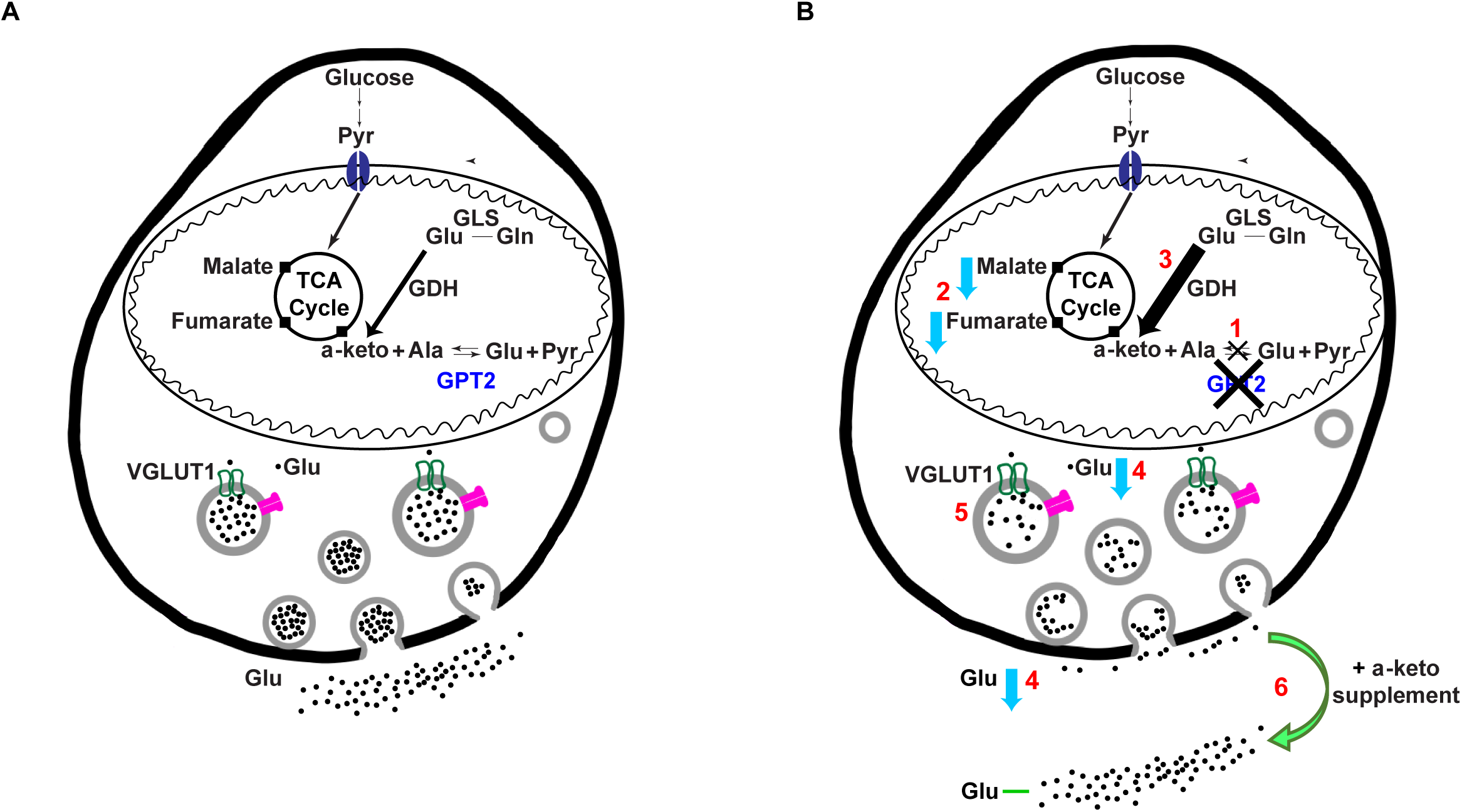
Loss of GPT2 leads to decreased pre-synaptic glutamate and synaptic transmission which is ameliorated by alpha-ketoglutarate supplementation. Based on the data, we propose the following model: **(1)** GPT2 is the main alanine aminotransferase and the main mediator of glutamine to alanine and glutamate to alanine interconversions (Fig. 1). **(2)** Loss of GPT2 leads to decreased TCA cycle intermediates in synaptosomes (Fig. 7). **(3)** Glutamate dehydrogenase (GDH) activity is increased (Fig. 6), likely to compensate for the deficit in the TCA cycle and to increase the flow of glutamate back into the TCA cycle. Double heavy nitrogen labeled glutamine pool is enriched (Fig. 6), likely to clear extra ammonia resulting from increased GDH activity, and concomitantly total glutamine levels are increased in *Gpt2*-null synaptosomes (Fig. 6). **(4)** We observe decreases in total glutamate in *Gpt2*-null synaptosomes as well as decreased glutamate levels released upon stimulation (Fig. 3 and 5). **(5)** We see increases in VGLUT1 protein levels in synaptosomes (Fig. S1) and the average size of synaptic vesicles in asymmetric glutamatergic spines in *Gpt2*-null CA1 stratum radiatum was also increased (Fig. 4), possibly to allow more glutamate entry into the synaptic vesicles. **(6)** Alpha-ketoglutarate supplementation in *Gpt2*-null synaptosomes increases released glutamate upon stimulation back to wild-type levels (Fig. 5).

There are anaplerotic pathways (i.e. resulting in a net yield of TCA cycle products), other than GDH and GPT2, that may support the TCA cycle in the absence of GPT2 such as pyruvate carboxylase (PC) and other aminotransferases that produce alpha-ketoglutarate from glutamate including branched-chain amino acid aminotransferases [56, 57]. All these processes likely contribute to sustain the TCA cycle in *Gpt2*-null brain as long as they are able; however, each process has certain complications. For example, PC exists mainly in astrocytes and utilizes ATP to make oxaloacetate [53, 58]. GDH is found both in neurons and astrocytes but depends on NADP availability [59]. Aminotransferases that degrade essential amino acids usually work in the direction of producing glutamate and not alpha-ketoglutarate [60]. In contrast, GPT2 is an equilibrium enzyme [61] and depends neither on ATP nor NADP availability, thereby conferring a unique anaplerotic ability to the cell. Levels of GPT2 and GPT1 protein have been shown to increase in models of oxidative phosphorylation deficits induced by mitochondrial DNA mutations [44] and disrupted mitochondrial fusion dynamics [43]. GPT2 may provide a non-redundant support of the TCA cycle that may enhance both energetics and biosynthesis to sustain cell growth.

Our data may suggest decreases in glutamate in pre-synaptic terminals and in synaptic vesicles (Fig. 8B, event 4). Evidence in support of this includes decreased mEPSCs amplitudes, and decreased release of glutamate from synaptosomes upon biochemical measurements. However, these data must be considered in context with other findings, such as increases in membrane resistance of CA1 pyramidal neurons. While none of our experiments alone can conclude that the glutamate pool is decreased specifically in neurons, the collection of our independent approaches supports this interpretation, including that there is also a reduction in overall glutamate levels in the cytosolic fraction of *Gpt2-*null forebrain (Fig. 3). It is difficult to bridge electrophysiological and biochemical data especially when it is hard to quantify glutamate levels in individual synaptic vesicles within synaptosomes, especially as the preparation involves using a hypoosmotic solution (ice-cold water). Structurally, we did not find changes in the number of synaptic vesicles per spine or number of spines per image area by electron microscopy. We do observe increases in synaptic vesicle area (Fig. 8B, event 5) and this could be due to compensatory increases in VGLUT1 protein expression as has been proposed to occur in similar situations [62].

In our studies, alpha-ketoglutarate rescues glutamate levels and augments the TCA cycle intermediates (Fig. 8B, event 6). It is yet unclear whether alpha-ketoglutarate exerts its effects through other transaminases or by blocking glutamate dehydrogenase, or both. Nevertheless, by a metabolic intervention, i.e. alpha-ketoglutarate supplementation, it was possible to correct both glutamate levels and alterations in the glutamine metabolism in *Gpt2-*null synaptosomes. Notably, decreases in alpha-ketoglutarate have been linked to decreased glutamatergic transmission [63]. Alanine on the other hand had minimal effect on glutamate levels and glutamine metabolism. It is possible that there was potentially a modest effect of alanine supplementation on glutamate levels in synaptosomes (p-value of 0.109) (Fig. 3). This is possible through the residual GPT1 in synaptosomes, however alanine failed to increase glutamate to wild-type levels and it failed to correct increased enrichment of double-labeled glutamine unlike alpha-ketoglutarate. Reduction in fraction of double-labeled glutamine most likely occurs through substrate inhibition of GDH by alpha-ketoglutarate [35].

Importantly, released GABA levels do not change in *Gpt2-*null synaptosomes. Glutamine and glutamate have been shown to be a major precursor for GABA whereas alpha-ketoglutarate is preferentially converted into glutamate and not GABA [31, 64, 65]. Our synaptosomes preparations contain not only glutamatergic but GABAergic synaptic terminals and presence of glutamate decarboxylase which irreversibly converts glutamate into GABA [66] possibly accounts for the unchanged GABA levels in *Gpt2-* null synaptosomes. We also observe increases in vesicular GABA transporter (VGAT) protein levels in *Gpt2-*null synaptosomes and this may indicate compensatory mechanisms acting on GABAergic synapses as well. Overall, we do not see changes in inhibitory synaptic transmission at the level of mIPSCs in hippocampal slices from *Gpt2-* null brain.

Synaptosomes, isolated synaptic terminals, are artificial organelles that may reflect both neuronal and astrocytic metabolism [20]. While we detected glutamine synthetase (GLUL), we do not observe a major common microtubule filament protein, GFAP in our synaptosome preparations. GLUL is found within astrocytic processes immediately next to excitatory spines by immuno-electron microscopy [67] and GLUL enzyme activity is demonstrated in synaptosome preps using Ficoll density gradient (similar to Percoll) despite reductions in glial membranes [23]. In addition, as our synaptosome preparations were obtained from a combination of cerebral cortex and hippocampus, it is not possible to correlate the findings in synaptosomes to one or the other of these specific brain regions; however, all of our electrophysiology data are conducted in hippocampus. Tripartite synapses (presynaptic terminals, postsynaptic spines and associated astrocyte components) exist only in approximately 50% of all spines in the stratum radiatum of the CA1 hippocampus [34]. Overall, the causal origin of the synaptic glutamate decreases has to be interpreted with caution. It is yet unclear whether GPT2 would have more profound effects on glutamatergic synapses that do not rely on astrocytic processes. Given our data, GPT2 may provide additional contributions to the TCA cycle at non-tripartite synapses *in vivo*.

Our prior work suggests that GPT2 is mainly neuronal and GPT1 is astrocytic. Compartmentation of metabolic enzymes and metabolites is a fundamental feature of the glutamine-glutamate cycle [10]. The consensus is that the glutamate pool is larger in neurons than glia [68] and given that GPT2 is an equilibrium enzyme [61], it is likely that GPT2 preferentially catalyzes the reaction in favor of alanine and alpha-ketoglutarate formation in neurons, thereby contributing to the TCA cycle metabolites in neurons. Moreover, there is evidence that alanine is preferentially released by neurons, to be subsequently taken in by astrocytes [69]. Most likely, the functions of the two glutamate pyruvate aminotransferase isozymes are not redundant and their individual importance arises from differences in expression in various cell types, subcellular localization and ranges of substrate metabolite pools in those cells.

GPT2 belongs to the protein family of pyridoxal phosphate (PLP)-dependent aspartate aminotransferase superfamily [70]. Aspartate aminotransferase 2 (GOT2) has been found to localize in both mitochondria and synaptic vesicles [71]. Localization to synaptic vesicles has been proposed as a mechanism to locally synthesize glutamate to be loaded into synaptic vesicles [72]. GPT2 does not appear in synaptic vesicles (Fig. 1D) and thus appears to modulate glutamate levels from within mitochondria; however, it is possible that the alpha-ketoglutarate rescue of glutamate release may function in part through GOT2 activity on synaptic vesicles [71, 72].

Processes related to glutamate metabolism may have major differences across species. Most notably, GPT1 seems to be absent in the human brain and GPT2 is the sole alanine aminotransferase detected [73, 74] whereas GPT1 expression is observed in murine brain tissue, particularly in astrocytes [75]. Another notable example is glutamate dehydrogenase. The *GDH* gene has seen a recent duplication event in humans and non-human primates, that is absent in rodents [76]. Therefore, caution must be taken when extrapolating interpretations of metabolic changes across species.

While it is possible that a peripheral mechanism may impose a long-standing effect on brain even after removed from the in vivo situation, there are several lines of evidence that support brain autonomous mechanisms based on: first, one of the promising results is that acute administration of alpha-ketoglutarate both augments glutamate release and brings down double heavy nitrogen labeling in glutamine in *Gpt2*-null synaptosomes. This effect is demonstrated ex vivo, outside any influence of peripheral tissues. Second, we have recently published that GPT2 is enriched in neurons as compared to astrocytes, and the conditional deletion of *Gpt2* in neuronal cells using *SynapsinI*-Cre driver results in a similar phenotype as the germline *Gpt2*-null whereas astrocyte-specific deletion of *Gpt2* using *Gfap*-Cre driver did not result in any overt phenotype [17]. However, a study of cell-specific deletions of *Gpt2* in the context of glutamate metabolism is warranted in the future.

In conclusion, here we report the contributions of GPT2 to glutamate metabolism and the functional consequences of GPT2 deficiency (Fig. 8). GPT2 loss results in impaired glutamatergic transmission in hippocampal slices. Decreased glutamate levels and altered glutamine metabolism in *Gpt2-*null synaptosomes can partly be alleviated by alpha-ketoglutarate. Compensatory mechanisms including upregulation of metabolic enzymes may act in lieu of GPT2 to sustain energetics and neurotransmitter availability. In sum, we find that GPT2 is a mitochondrial enzyme that links TCA cycle metabolism with glutamate metabolism and thereby influences glutamatergic synaptic transmission. Our work identifies GPT2 as a mitochondrial enzyme that modulates synaptic glutamate, and alpha-ketoglutarate supplementation as a potential therapeutic in human patients with GPT2 deficiency.

## Availability of data and materials

Further data and reagents used in this manuscript can be shared upon request.

## Authors’ contributions

Conceptualization O.B., J.A.K. and E.M.M.; O.B. conducted all experiments; S.M.D. contributed to studies involving mass spectrometry for isotope tracing; Data Curation and Visualization O.B.; Experimental Design and Data Analysis O.B., S.M.D., J.A.K. and E.M.M.; Formal Analysis and Writing – Original Draft, O.B. and E.M.M.; Writing – Review & Editing, O.B., S.M.D., J.A.K. and E.M.M; Supervision, Project Administration, and Funding Acquisition, J.A.K. and E.M.M.

## Acknowledgements

We would like to thank Paula Weston (Brown University, Molecular Pathology Core) and Maria Ericsson (Harvard Medical School) for their help with electron microscopy.

## Supporting Information

**Figure S1.**
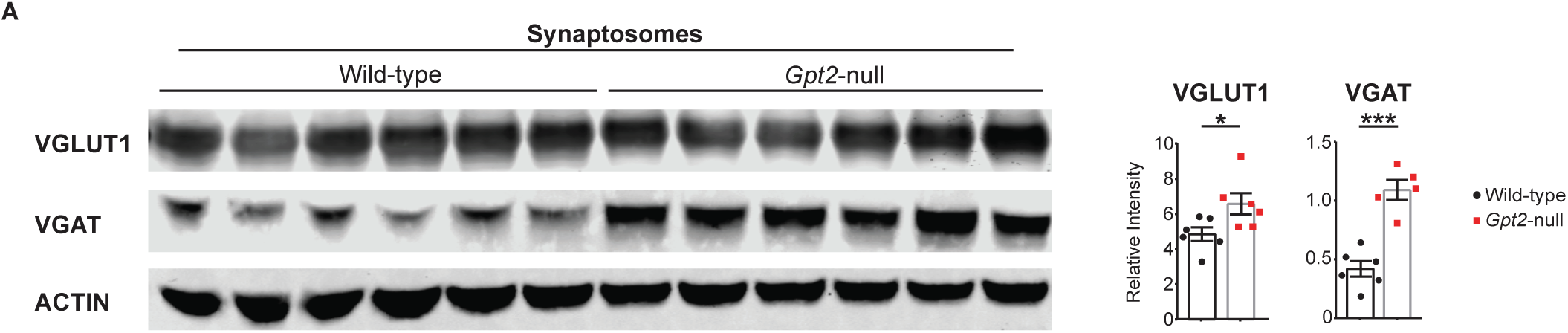
VGLUT1 and VGAT protein levels are increased in Gpt2-null synaptosomes. **A.** Western blotting of wild-type and Gpt2-null synaptosome protein lysates collected at P18. Each dot represents a different protein lysate sample from wild-type (black) or Gpt2-null (red) mice. Each protein band is normalized to its corresponding intensity of the actin band. Wild-type vs. Gpt2-null: *P (VGLUT1, vesicular glutamate transporter 1) = 0.038; ***P (VGAT, vesicular GABA transporter) = 0.0001.

**Figure S2.**
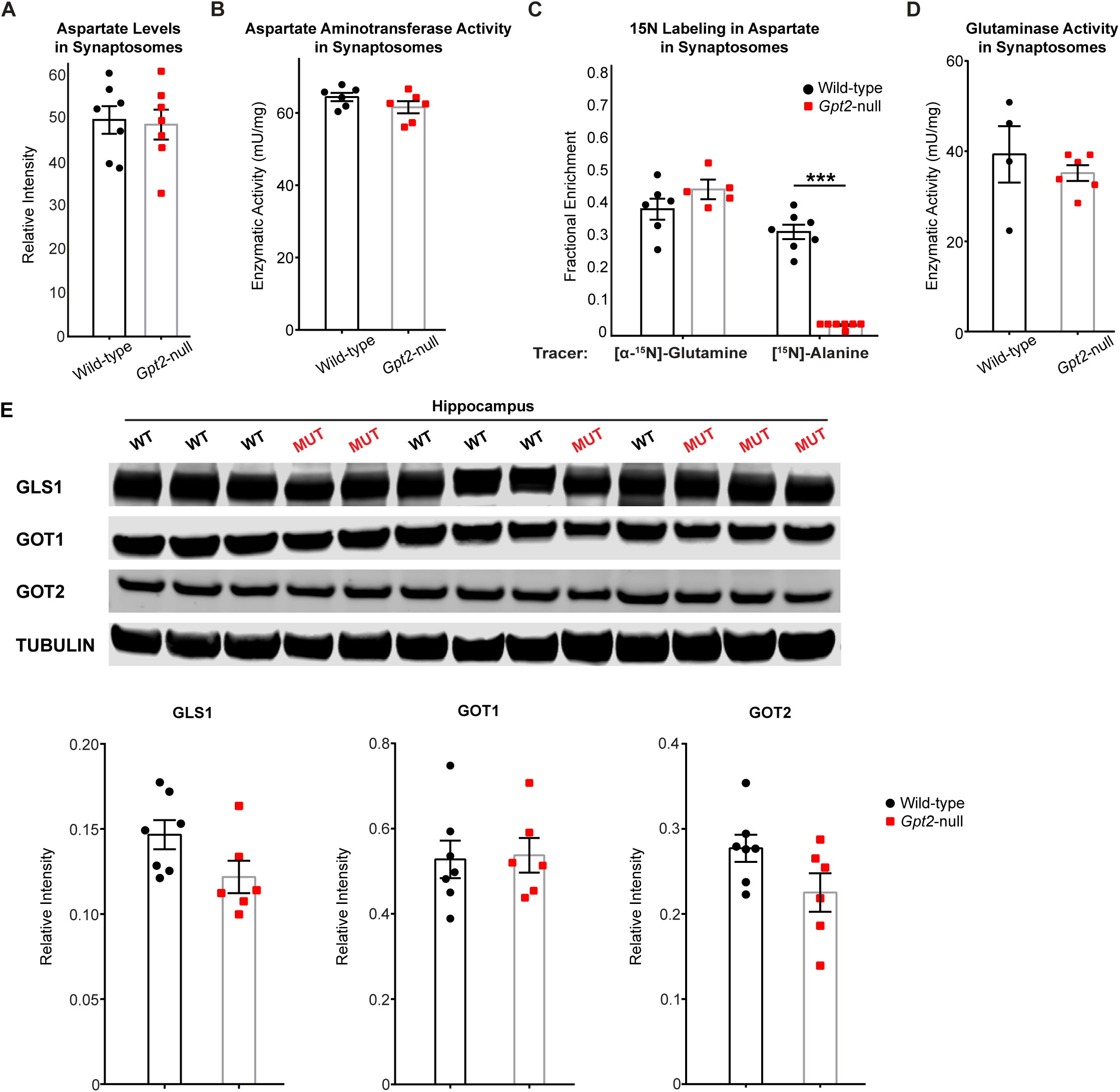
Aspartate aminotransferase and glutaminase enzyme activities in *Gpt2*-null synaptosomes. **A.** Aspartate levels in *Gpt2*-null synaptosomes. Each dot represents a different synaptosome sample obtained from wild-type (black) or *Gpt2*-null (red) mice at P18. **B.** Aspartate aminotransferase enzyme activity in wild-type and *Gpt2*-null synaptosomes. Each dot represents a different synaptosome sample obtained from wild-type (black) or *Gpt2*-null (red) mice at P18. **C.** 15N (heavy nitrogen) labeling of aspartate in *Gpt2*-null synaptosomes using [α-15N]-glutamine or [α-15N]-alanine as tracers. Each dot represents a different synaptosome sample obtained from wild-type (black) or *Gpt2*-null (red) mice at P18. \*\*\**P* <0.0001. **D.** Glutaminase enzyme activity in wild-type and *Gpt2*-null synaptosomes. Each dot represents a different synaptosome sample obtained from wild-type (black) or *Gpt2*-null (red) mice at P18. **E.** Western blotting for aspartate aminotransferase 1 (GOT1), aspartate aminotransferase 2 (GOT2) and glutaminase (GLS1) in wild-type and *Gpt2*-null hippocampus protein lysates. Each dot represents a different synaptosome sample obtained from wild-type (black) or *Gpt2*-null (red) mice at P18. Each protein band is normalized to its corresponding intensity of the tubulin band.

**Figure S3.**
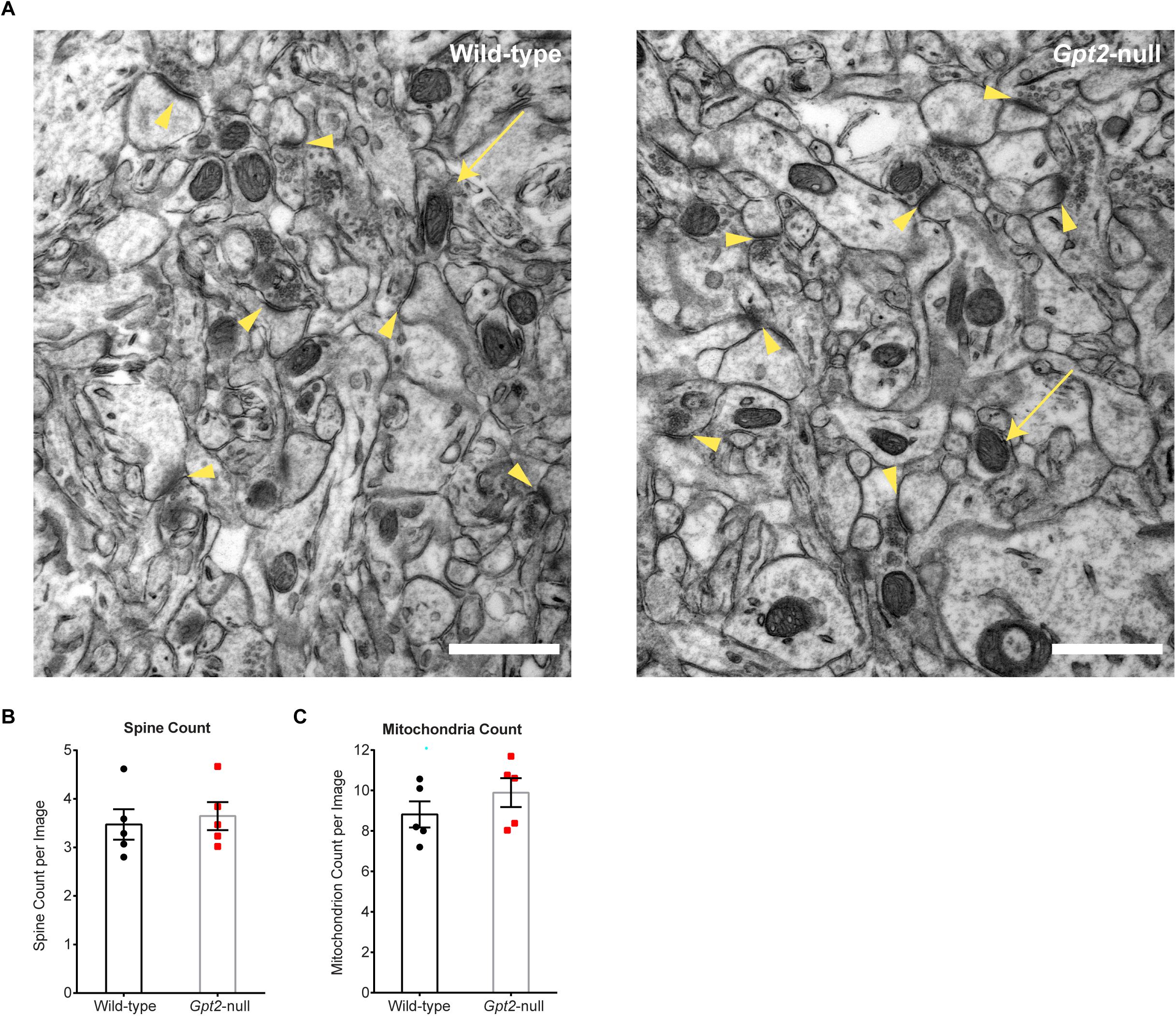
Asymmetric spine and mitochondria counts are unchanged in electron micrographs of CA1 stratum radiatum of *Gpt2*-null hippocampus. **A.** Representative electron micrographs of wild-type (left) and *Gpt2*-null (right) CA1 stratum radiatum of the hippocampus at P18. Orange arrowheads point to the asymmetric spines, arrows point to the mitochondria. Scale bar: 1 µm. **B.** Quantification of spines per image (21000X magnification). Each dot represents the averaged number of spines obtained from one animal. More than 25 micrographs from each animal were analyzed. **C.** Quantification of mitochondria per image (21000X magnification). Each dot represents the averaged number of mitochondria obtained from one animal. More than 25 micrographs from each animal were analyzed.

**Figure S4.**
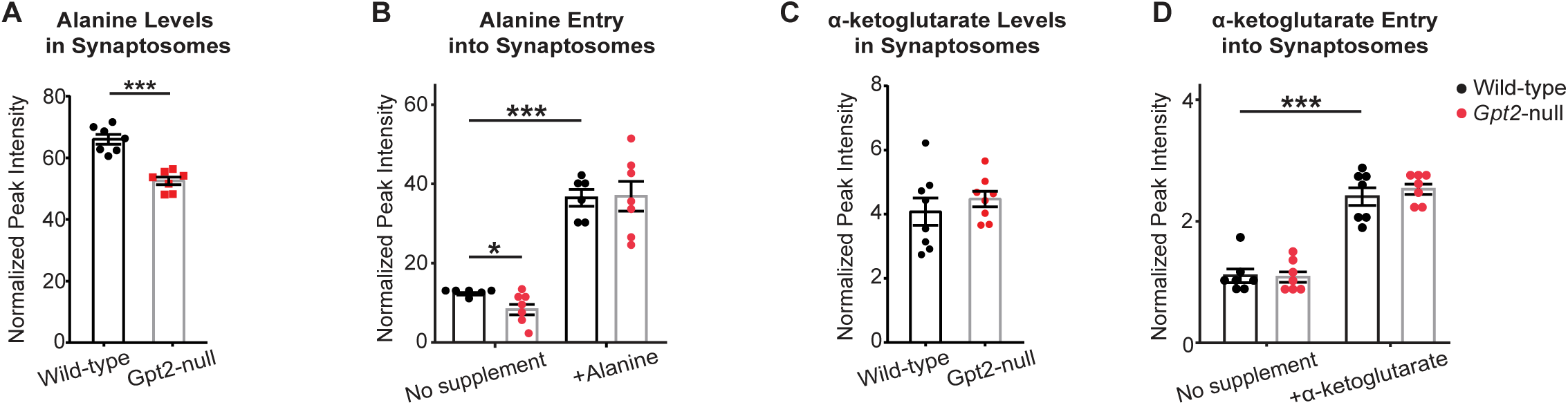
Alanine and alpha-ketoglutarate readily enter synaptosomes. **A.** Alanine levels are reduced in *Gpt2*-null synaptosomes. Each dot represents a different synaptosome sample obtained from wild-type (black) or *Gpt2*-null (red) mice at P18. \*\*\**P* <0.0001. **B.** Alanine can readily enter the synaptosomes. Each dot represents a different synaptosome sample obtained from wild-type (black) or *Gpt2*-null (red) mice at P18. No supplement: wild-type vs. *Gpt2*-null: \**P* = 0.02. Wild-type no supplement vs. wild-type +alanine: \*\*\**P* <0.0001. +alanine: wild-type vs. *Gpt2*-null: *P* = 0.94. **C.** Alpha-ketoglutarate levels are not changed in *Gpt2*-null synaptosomes. Each dot represents a different synaptosome sample obtained from wild-type (black) or *Gpt2*-null (red) mice at P18. *P* = 0.43. **D.** Alpha-ketoglutarate can readily enter the synaptosomes. Each dot represents a different synaptosome sample obtained from wild-type (black) or *Gpt2*-null (red) mice at P18. Wild-type no supplement vs. wild-type +alpha-ketoglutarate: \*\*\**P* <0.0001.

**Figure S5.**
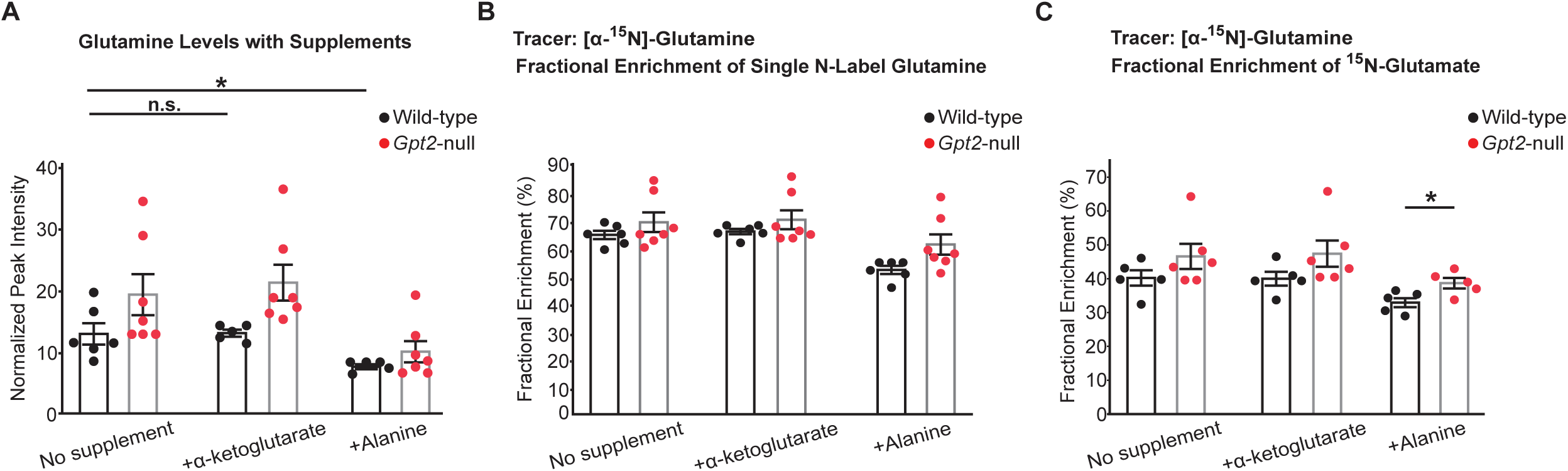
Glutamine entry and nitrogen labeling of glutamine and glutamate in *Gpt2*-null synaptosomes. **A.** Entry of glutamine is unaffected by alpha-ketoglutarate. Each dot represents synaptosome samples obtained from different wild-type (black) or *Gpt2*-null (red) mice at P18. Wild-type no supplement vs. +alanine: \**P* = 0.02; Wild-type no supplement vs. +alpha-ketoglutarate: *P* = 0.95. **B.** Exogenous amine labeled glutamine labels the majority of glutamine pool in synaptosomes. Each dot represents synaptosome samples obtained from different wild-type (black) or *Gpt2*-null (red) mice at P18. Wild-type vs. *Gpt2*-null no supplement: *P* = 0.29; +alanine: *P* = 0.0513; +alpha-ketoglutarate: *P* = 0.29. **C.** Fractional enrichment of labeled glutamate with the amine heavy nitrogen labeled glutamine precursor. Each dot represents synaptosome samples obtained from different wild-type (black) or *Gpt2*-null (red) mice at P18. Wild-type vs. *Gpt2*-null no supplement: *P* = 0.2; +alanine: \**P* = 0.023; +alpha-ketoglutarate: *P* = 0.15.

**Figure S6.**
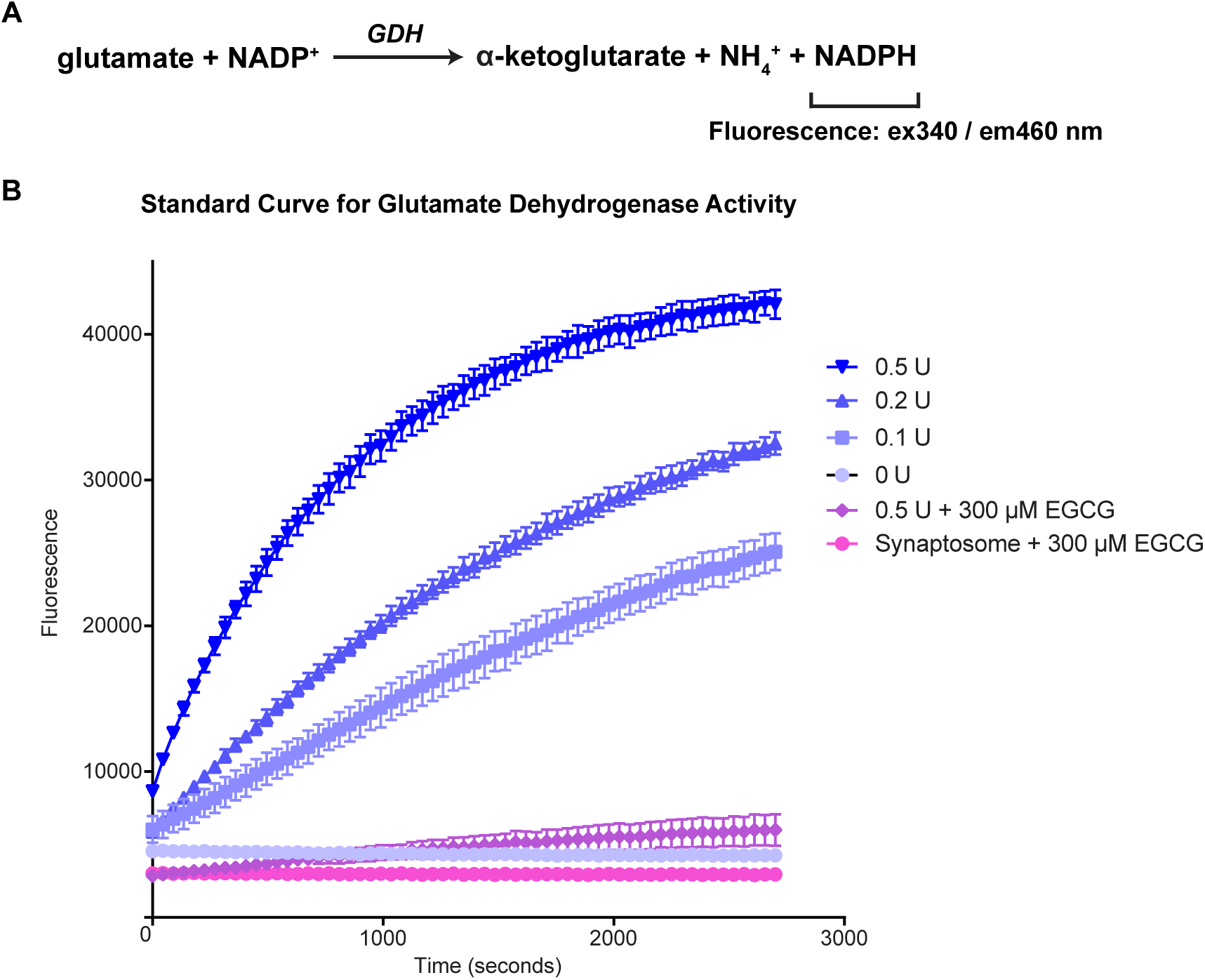
Confirmation of glutamate dehydrogenase enzyme activity in synaptosomes. **A.** Chemical reaction catalyzed by glutamate dehydrogenase (GDH). The enzyme activity assay detects fluorescence with excitation and emission wavelengths of 340 and 460 nm, respectively. **B.** Standard curve used to determine GDH enzymatic activity in synaptosomes. 300 µM Epigallocatechin gallate (EGCG) is used as a glutamate dehydrogenase inhibitor to validate the assay [77].

## References

1. Fonnum, F., Glutamate: a neurotransmitter in mammalian brain. J Neurochem, 1984. 42(1): p. 1–11.

2. Villa, K.L. and E. Nedivi, Excitatory and Inhibitory Synaptic Placement and Functional Implications, in Dendrites. 2016. p. 467–487.

3. Kovacevic, Z., The pathway of glutamine and glutamate oxidation in isolated mitochondria from mammalian cells. Biochem J, 1971. 125(3): p. 757–63.

4. Westergaard, N., U. Sonnewald, and A. Schousboe, Metabolic trafficking between neurons and astrocytes: the glutamate/glutamine cycle revisited. Dev Neurosci, 1995. 17(4): p. 203–11.

5. Kurmi, K. and M.C. Haigis, Nitrogen Metabolism in Cancer and Immunity. Trends Cell Biol, 2020. 30(5): p. 408–424.

6. Cooper, A.J. and T.M. Jeitner, Central Role of Glutamate Metabolism in the Maintenance of Nitrogen Homeostasis in Normal and Hyperammonemic Brain. Biomolecules, 2016. 6(2).

7. Young, V.R. and A.M. Ajami, Glutamate: an amino acid of particular distinction. J Nutr, 2000. 130(4S Suppl): p. 892S–900S.

8. Sedlak, T.W., et al., The glutathione cycle shapes synaptic glutamate activity. Proc Natl Acad Sci U S A, 2019. 116(7): p. 2701–2706.

9. Sonnewald, U. and M. McKenna, Metabolic compartmentation in cortical synaptosomes: influence of glucose and preferential incorporation of endogenous glutamate into GABA. Neurochem Res, 2002. 27(1-2): p. 43–50.

10. Sonnewald, U. and A. Schousboe, Introduction to the Glutamate-Glutamine Cycle. Adv Neurobiol, 2016. 13: p. 1–7.

11. Ouyang, Q., et al., Mutations in mitochondrial enzyme GPT2 cause metabolic dysfunction and neurological disease with developmental and progressive features. Proc Natl Acad Sci U S A, 2016. 113(38): p. E5598–607.

12. Lindblom, P., et al., Isoforms of alanine aminotransferases in human tissues and serum--differential tissue expression using novel antibodies. Arch Biochem Biophys, 2007. 466(1): p. 66–77.

13. Smith, B., et al., Addiction to Coupling of the Warburg Effect with Glutamine Catabolism in Cancer Cells. Cell Rep, 2016. 17(3): p. 821–836.

14. Kim, M., et al., Mitochondrial GPT2 plays a pivotal role in metabolic adaptation to the perturbation of mitochondrial glutamine metabolism. Oncogene, 2019. 38(24): p. 4729–4738.

15. Jin, L., et al., Glutamate dehydrogenase 1 signals through antioxidant glutathione peroxidase 1 to regulate redox homeostasis and tumor growth. Cancer Cell, 2015. 27(2): p. 257–70.

16. McCommis, K.S., et al., Loss of Mitochondrial Pyruvate Carrier 2 in the Liver Leads to Defects in Gluconeogenesis and Compensation via Pyruvate-Alanine Cycling. Cell Metab, 2015. 22(4): p. 682–94.

17. Baytas, O., S.M. Davidson, R.J. DeBerardinis, and E.M. Morrow, Mitochondrial enzyme GPT2 regulates metabolic mechanisms required for neuron growth and motor function in vivo. Hum Mol Genet, 2021.

18. Council, N.R., Guide for the Care and Use of Laboratory Animals: Eighth Edition. 2011, Washington, DC: The National Academies Press. 246.

19. Sims, N.R. and M.F. Anderson, Isolation of mitochondria from rat brain using Percoll density gradient centrifugation. Nat Protoc, 2008. 3(7): p. 1228–39.

20. Dunkley, P.R., P.E. Jarvie, and P.J. Robinson, A rapid Percoll gradient procedure for preparation of synaptosomes. Nat Protoc, 2008. 3(11): p. 1718–28.

21. Brown, T.E., A.M. Chirila, B.R. Schrank, and J.A. Kauer, Loss of interneuron LTD and attenuated pyramidal cell LTP in Trpv1 and Trpv3 KO mice. Hippocampus, 2013. 23(8): p. 662–71.

22. Gulyassy, P., et al., Proteomic comparison of different synaptosome preparation procedures. Amino Acids, 2020. 52(11-12): p. 1529–1543.

23. Dennis, S.C., J.C.K. Lai, and J.B. Clark, The distribution of glutamine synthetase in subcellular fractions of rat brain. Brain Research, 1980. 197(2): p. 469–475.

24. Byun, Y.G. and W.S. Chung, A Novel In Vitro Live-imaging Assay of Astrocyte-mediated Phagocytosis Using pH Indicator-conjugated Synaptosomes. J Vis Exp, 2018(132).

25. Rimmele, T.S. and P.A. Rosenberg, GLT-1: The elusive presynaptic glutamate transporter. Neurochem Int, 2016. 98: p. 19–28.

26. Ropert, N., R. Miles, and H. Korn, Characteristics of miniature inhibitory postsynaptic currents in CA1 pyramidal neurones of rat hippocampus. J Physiol, 1990. 428: p. 707–22.

27. Fatt, P. and B. Katz, Spontaneous Subthreshold Activity At Motor Nerve Endings. J Physiol, 1952.

28. Benuck, M. and A. Lajtha, Aminotransferase activity in brain. Int Rev Neurobiol, 1975. 17: p. 85–129.

29. Shank, R.P., W.J. Baldy, and C.W. Ash, Glutamine and 2-oxoglutarate as metabolic precursors of the transmitter pools of glutamate and GABA: correlation of regional uptake by rat brain synaptosomes. Neurochem Res, 1989. 14(4): p. 371–6.

30. Peng, L.A., A. Schousboe, and L. Hertz, Utilization of alpha-ketoglutarate as a precursor for transmitter glutamate in cultured cerebellar granule cells. Neurochem Res, 1991. 16(1): p. 29–34.

31. Peng, L., et al., Utilization of glutamine and of TCA cycle constituents as precursors for transmitter glutamate and GABA. Dev Neurosci, 1993. 15(3-5): p. 367–77.

32. Shank, R.P. and D.J. Bennett, 2-Oxoglutarate transport: a potential mechanism for regulating glutamate and tricarboxylic acid cycle intermediates in neurons. Neurochem Res, 1993. 18(4): p. 401–10.

33. Baytas, O., S.M. Davidson, R.J. DeBerardinis, and E.M. Morrow, Mitochondrial enzyme GPT2 regulates metabolic mechanisms required for neuron growth and motor function in vivo. Hum Mol Genet, 2022. 31(4): p. 587–603.

34. Ventura, R. and K.M. Harris, Three-Dimensional Relationships between Hippocampal Synapses and Astrocytes. The Journal of Neuroscience, 1999. 19(16): p. 6897–6906.

35. Engel, P.C. and S.S. Chen, A product-inhibition study of bovine liver glutamate dehydrogenase. Biochem J, 1975. 151(2): p. 305–18.

36. Leke, R. and A. Schousboe, The Glutamine Transporters and Their Role in the Glutamate/GABA-Glutamine Cycle. Adv Neurobiol, 2016. 13: p. 223–257.

37. Erecinska, M. and I.A. Silver, Metabolism and role of glutamate in mammalian brain. Prog Neurobiol, 1990. 35(4): p. 245–96.

38. Owen, O.E., S.C. Kalhan, and R.W. Hanson, The key role of anaplerosis and cataplerosis for citric acid cycle function. J Biol Chem, 2002. 277(34): p. 30409–12.

39. Haslam, R.J. and H.A. Krebs, The Metabolism of Glutamate in Homogenates and Slices of Brain Cortex. Biochem J, 1963. 88: p. 566–78.

40. Schousboe, A., N. Westergaard, and L. Hertz, Neuronal-astrocytic interactions in glutamate metabolism. Biochem Soc Trans, 1993. 21(1): p. 49–53.

41. Yudkoff, M., et al., Brain glutamate metabolism: neuronal-astroglial relationships. Dev Neurosci, 1993. 15(3-5): p. 343–50.

42. Anderson, C.M. and R.A. Swanson, Astrocyte glutamate transport: Review of properties, regulation, and physiological functions. Glia, 2000. 32(1): p. 1–14.

43. Motori, E., et al., Neuronal metabolic rewiring promotes resilience to neurodegeneration caused by mitochondrial dysfunction. Sci Adv, 2020. 6(35): p. eaba8271.

44. Chen, Q., et al., Rewiring of Glutamine Metabolism Is a Bioenergetic Adaptation of Human Cells with Mitochondrial DNA Mutations. Cell Metab, 2018. 27(5): p. 1007–1025 e5.

45. Yang, C., et al., Glutamine oxidation maintains the TCA cycle and cell survival during impaired mitochondrial pyruvate transport. Mol Cell, 2014. 56(3): p. 414–24.

46. Alkan, H.F., et al., Cytosolic Aspartate Availability Determines Cell Survival When Glutamine Is Limiting. Cell Metab, 2018. 28(5): p. 706–720 e6.

47. Hohnholt, M.C., et al., Glutamate dehydrogenase is essential to sustain neuronal oxidative energy metabolism during stimulation. J Cereb Blood Flow Metab, 2018. 38(10): p. 1754–1768.

48. Bowman, C.E., L. Zhao, T. Hartung, and M.J. Wolfgang, Requirement for the Mitochondrial Pyruvate Carrier in Mammalian Development Revealed by a Hypomorphic Allelic Series. Mol Cell Biol, 2016. 36(15): p. 2089–104.

49. Kanehisa, M., et al., KEGG: integrating viruses and cellular organisms. Nucleic Acids Res, 2020.

50. Yoo, H.C., Y.C. Yu, Y. Sung, and J.M. Han, Glutamine reliance in cell metabolism. Exp Mol Med, 2020. 52(9): p. 1496–1516.

51. Brekke, E., et al., Anaplerosis for Glutamate Synthesis in the Neonate and in Adulthood. Adv Neurobiol, 2016. 13: p. 43–58.

52. McKenna, M.C., The glutamate-glutamine cycle is not stoichiometric: fates of glutamate in brain. J Neurosci Res, 2007. 85(15): p. 3347–58.

53. Hertz, L., L. Peng, and G.A. Dienel, Energy metabolism in astrocytes: high rate of oxidative metabolism and spatiotemporal dependence on glycolysis/glycogenolysis. J Cereb Blood Flow Metab, 2007. 27(2): p. 219–49.

54. McKenna, M.C., J.H. Stevenson, X. Huang, and I.B. Hopkins, Differential distribution of the enzymes glutamate dehydrogenase and aspartate aminotransferase in cortical synaptic mitochondria contributes to metabolic compartmentation in cortical synaptic terminals. Neurochem Int, 2000. 37(2-3): p. 229–41.

55. Westergaard, N., J. Drejer, A. Schousboe, and U. Sonnewald, Evaluation of the importance of transamination versus deamination in astrocytic metabolism of [U-13C] glutamate. Glia, 1996. 17(2): p. 160–168.

56. Morken, T.S., et al., Neuron-astrocyte interactions, pyruvate carboxylation and the pentose phosphate pathway in the neonatal rat brain. Neurochem Res, 2014. 39(3): p. 556–69.

57. Hutson, S., Structure and function of branched chain aminotransferases. 2001. p. 175–206.

58. Yu, A.C., J. Drejer, L. Hertz, and A. Schousboe, Pyruvate carboxylase activity in primary cultures of astrocytes and neurons. J Neurochem, 1983. 41(5): p. 1484–7.

59. Aoki, C., et al., Regional distribution of astrocytes with intense immunoreactivity for glutamate dehydrogenase in rat brain: implications for neuron-glia interactions in glutamate transmission. The Journal of Neuroscience, 1987. 7(7): p. 2214–2231.

60. Conway, M.E. and S.M. Hutson, BCAA Metabolism and NH3 Homeostasis. Adv Neurobiol, 2016. 13: p. 99–132.

61. Krebs, H.A., Equilibria in transamination systems. Biochem J, 1953. 54(1): p. 82–6.

62. Daniels, R.W., et al., Increased expression of the Drosophila vesicular glutamate transporter leads to excess glutamate release and a compensatory decrease in quantal content. J Neurosci, 2004. 24(46): p. 10466–74.

63. Ugur, B., et al., The Krebs Cycle Enzyme Isocitrate Dehydrogenase 3A Couples Mitochondrial Metabolism to Synaptic Transmission. Cell Rep, 2017. 21(13): p. 3794–3806.

64. Shank, R.P. and G.L. Campbell, Glutamine and alpha-ketoglutarate uptake and metabolism by nerve terminal enriched material from mouse cerebellum. Neurochem Res, 1982. 7(5): p. 601–16.

65. Shank, R.P. and G.L. Campbell, Alpha-ketoglutarate and malate uptake and metabolism by synaptosomes: further evidence for an astrocyte-to-neuron metabolic shuttle. J Neurochem, 1984. 42(4): p. 1153–61.

66. Roberts, E. and E. Eidelberg, Metabolic and Neurophysiological Roles of γ-Aminobutyric Acid. 1960. p. 279–332.

67. Derouiche, A. and M. Frotscher, Astroglial processes around identified glutamatergic synapses contain glutamine synthetase: evidence for transmitter degradation. Brain Res, 1991. 552(2): p. 346–50.

68. Erecinska, M. and I. Silver, Metabolism and role of glutamate in mammalian brain. Progress in Neurobiology, 1990. 35(4): p. 245–296.

69. Waagepetersen, H.S., U. Sonnewald, O.M. Larsson, and A. Schousboe, A possible role of alanine for ammonia transfer between astrocytes and glutamatergic neurons. J Neurochem, 2000. 75(2): p. 471–9.

70. Yang, R.Z., et al., cDNA cloning, genomic structure, chromosomal mapping, and functional expression of a novel human alanine aminotransferase. Genomics, 2002. 79(3): p. 445–50.

71. Takeda, K. and T. Ueda, Effective Mechanism for Synthesis of Neurotransmitter Glutamate and its Loading into Synaptic Vesicles. Neurochem Res, 2017. 42(1): p. 64–76.

72. Takeda, K., A. Ishida, K. Takahashi, and T. Ueda, Synaptic vesicles are capable of synthesizing the VGLUT substrate glutamate from alpha-ketoglutarate for vesicular loading. J Neurochem, 2012. 121(2): p. 184–96.

73. Kang, H.J., et al., Spatio-temporal transcriptome of the human brain. Nature, 2011. 478(7370): p. 483–9.

74. Hodge, R.D., et al., Conserved cell types with divergent features in human versus mouse cortex. Nature, 2019. 573(7772): p. 61–68.

75. Zhang, Y., et al., Purification and Characterization of Progenitor and Mature Human Astrocytes Reveals Transcriptional and Functional Differences with Mouse. Neuron, 2016. 89(1): p. 37–53.

76. Li, Q., et al., Mice carrying a human GLUD2 gene recapitulate aspects of human transcriptome and metabolome development. Proc Natl Acad Sci U S A, 2016. 113(19): p. 5358–63.

77. Li, C., et al., Green tea polyphenols modulate insulin secretion by inhibiting glutamate dehydrogenase. J Biol Chem, 2006. 281(15): p. 10214–21.

